# Global transcriptional rewiring provides recurrent evolutionary routes to insect–*Escherichia coli* mutualism

**DOI:** 10.64898/2026.08.13.744627

**Authors:** Yayun Wang, Ryuga Sugiyama, Yudai Nishide, Tomonari Nozaki, Masaki Mizutani, Ryutaro Suzuki, Minoru Moriyama, Ryuichi Koga, Takema Fukatsu

## Abstract

Microbial symbionts often evolve specialized functions that compensate for host nutritional deficiencies, yet the evolutionary routes by which such mutualisms arise remain poorly understood. Here, using an experimental symbiosis between the stinkbug *Plautia stali* and *Escherichia coli*, we investigated the early evolution of insect–bacterium mutualism. Among 144 independently evolved bacterial lineages, six acquired the ability to improve host performance. Four lineages carried independent disruptive mutations in *cyaA*, disabling carbon catabolite repression and thereby increasing the essential amino acid tryptophan. The remaining two lineages evolved through nonsynonymous mutations in *rpoB* or *rpoD*, encoding core components of the transcriptional machinery. Transcriptomic analyses revealed extensive regulatory rewiring in both mutants, including marked upregulation of biosynthetic pathways for branched-chain essential amino acids. Our results identify global transcriptional regulatory systems as major evolutionary targets for the emergence of mutualism and demonstrate how large-effect regulatory mutations can rapidly generate host-beneficial traits. (146 < 150 words)

## Introduction

Microbial symbionts are widespread across animals, plants, fungi and protists, and contribute fundamentally to host diversification, ecology and environmental adaptation^1–5^. Long-term host–symbiont coevolution often produces extreme bacterial specialization, including genome reduction, extensive gene loss, uncultivability, and metabolic streamlining towards host-beneficial functions such as the provisioning of vitamins, essential amino acids and digestive enzymes^6–9^. Yet these highly specialized symbionts ultimately descend from non-symbiotic, often free-living ancestors. A central unresolved question is therefore what genetic changes first convert an ordinary bacterium into a beneficial partner and whether this transition follows some specific evolutionary trajectories. Experimental evolution provides a direct means to identify these early causal changes^10–13^.

Recently, the stinkbug *Plautia stali* and *Escherichia coli* have been developed as an experimental system for studying the evolution of mutualism^13–18^. *P. stali* normally harbors an essential *Pantoea* symbiont in a specialized midgut organ^19–22^, whereas *E. coli* is a mammalian gut bacterium with no prior evolutionary association with this insect. Nevertheless, serial passage of a hypermutator *E. coli* strain through symbiont-deprived *P. stali* yielded bacterial lineages that improved host adult emergence and body coloration within months^14^. In two independently evolved mutualistic strains, disruptive mutations in *cyaA* or *crp* convergently disabled carbon catabolite repression (CCR), the bacterial global transcriptional regulatory system for switching metabolism, and thereby caused the host-beneficial phenotypes^14^. Subsequent functional analyses identified downstream metabolic routes by which these single regulatory mutations can benefit the host: disruption of *tnaA*, encoding tryptophanase, increases an essential amino acid tryptophan and reduces toxic indole production^15^. Also, disruption of *metJ*, encoding a methionine biosynthesis repressor, was shown to increase another essential amino acid, methionine, and thereby contribute to host fitness^16^. These findings established that *E. coli* can rapidly acquire host-beneficial functions, but the limited number of evolved lineages left unresolved whether CCR disruption represents a dominant and reproducible route to mutualism or only one of many accessible trajectories.

To determine the breadth and repeatability of evolutionary routes to mutualism, we conducted large-scale experimental evolution using 144 independently maintained *P. stali*–*E. coli* lineages. Six lineages evolved to improve host performance. Four lineages independently acquired mutations in *cyaA*, demonstrating recurrent evolution through disruption of CCR. The remaining two lineages carried host-beneficial mutations in *rpoB* and *rpoD*, which encode core components of the transcriptional machinery. These mutations caused extensive transcriptional rewiring, including increased expression of biosynthesis pathways for essential branched-chain amino acids. Thus, independent evolutionary trajectories converged at a higher functional level on global transcriptional regulation, identifying regulatory network reprogramming as a major early route to insect–bacterium mutualism.

## Results

### Large-scale experimental evolution identifies candidate mutualistic lineages

We established 144 independent *P. stali*–*E. coli* evolutionary lineages under two transmission regimes (Fig. 1). The vertical-transmission experiment comprised 58 hypermutator Δ*mutS* lineages (Vm01–Vm58) and 56 control Δ*intS* lineages (Vi01–Vi56), whereas the artificial-inoculation experiment comprised 15 Δ*mutS* lineages (Am01–Am15) and 15 Δ*intS* lineages (Ai01–Ai15). Adult emergence and body coloration were monitored across generations. For the initial screen, we retained lineages that persisted stably for multiple generations and reached adult emergence rates above 30% multiple times for vertical-transmission lineages and above 45% multiple times for artificial-inoculation lineages, whereas a few lineages were additionally retained on account of color and size of the infected insects (Fig. 2). This deliberately permissive criterion identified 21 candidates: 17 vertical-transmission Δ*mutS* lineages, one vertical-transmission Δ*intS* lineage and three artificial-inoculation Δ*mutS* lineages (Table S1).

**Fig. 1.**
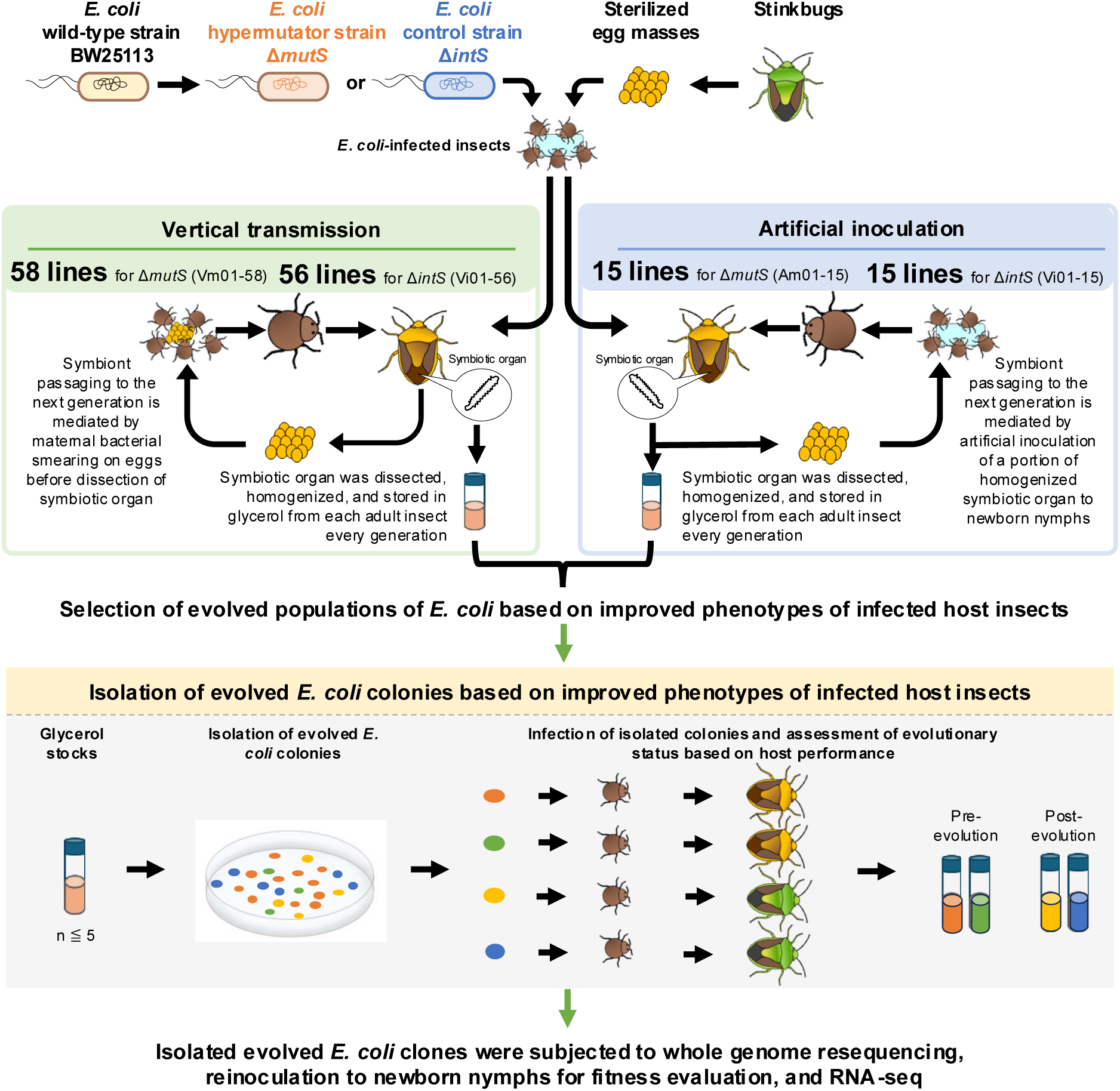
Large-scale evolutionary experiments using the *P. stali*–*E. coli* artificial symbiotic system. In total, 114 evolutionary *E. coli* lineages were passaged through host generations by vertical transmission, whereas 30 evolutionary *E. coli* lineages were passaged by artificial inoculation.

**Fig. 2.**
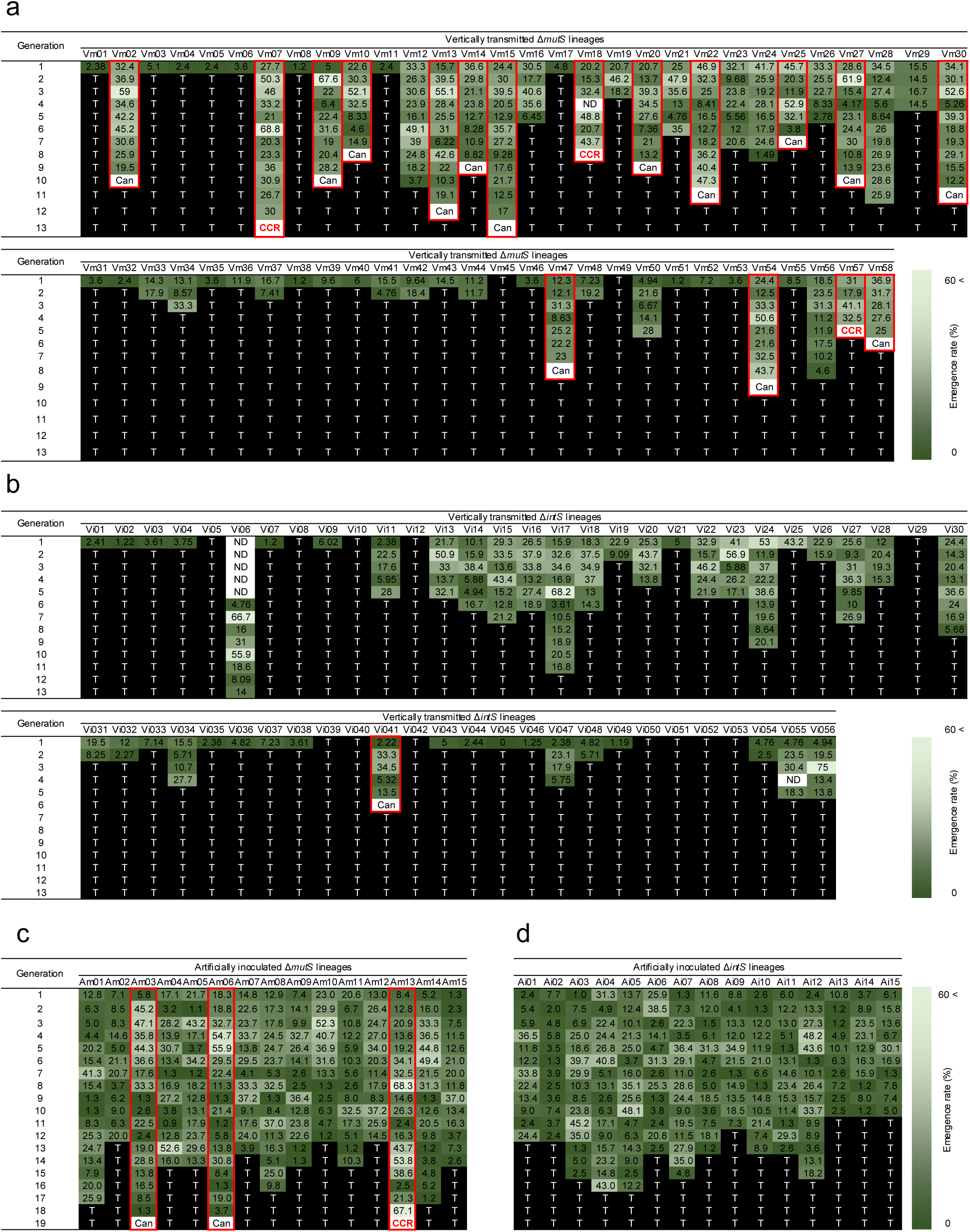
Initial screening of evolved *E. coli* lineages based on host adult emergence. (**a–d**) Heatmaps showing adult emergence rates across host generations for (**a**) 58 Δ*mutS* lineages (Vm01–Vm58) passaged by vertical transmission, (**b**) 56 Δ*intS* lineages (Vi01–Vi56) passaged by vertical transmission, (**c**) 15 Δ*mutS* lineages (Am01–Am15) passaged by artificial inoculation and (**d**) 15 Δ*intS* lineages (Ai01–Ai15) passaged by artificial inoculation. Red boxes indicate candidate lineages selected for further analysis. CCR, candidate lineage carrying a mutation in the CCR-associated gene *cyaA*; Can, candidate lineage lacking mutations in *cyaA* or *crp*; ND, no data; T, lineage extinction or termination of passaging.

### Recurrent *cyaA* mutations underlie independently evolved host-beneficial phenotypes

Targeted sequencing of *cyaA* and *crp* in the 21 candidate lineages identified *cyaA* mutations in four independent lineages: a frameshift insertion in Vm07, a nonsense mutation in Vm18, and nonsynonymous substitutions in Vm57 and Am13 (Table S1). Whole-genome resequencing of twelve Vm07 clones, eight Vm18 clones and fifteen Vm57 clones isolated across host generations confirmed evolutionary emergence of these *cyaA* variants (Tables S2 and S3). Reinoculation of archived Vm07, Vm18 and Vm57 populations into symbiont-free *P. stali* significantly increased adult emergence within one to four host generations (Fig. S1a–c). Adult coloration also improved in Vm07 and Vm18, but not in Vm57 (Fig. S1d–f). Thus, four independent lineages acquired *cyaA* mutations, and three tested lineages evolved reproducible host-beneficial phenotypes, supporting CCR disruption as a recurrent route to *P. stali*–*E. coli* mutualism.

### Evolution of mutualistic *E. coli* lineages without CCR disruption

To identify evolutionary routes independent of CCR disruption, we reinoculated frozen stocks from the 17 remaining candidate lineages lacking *cyaA* or *crp* mutations: Vm02, Vm09, Vm10, Vm13, Vm15, Vm16, Vm20, Vm22, Vm25, Vm27, Vm30, Vm47, Vm54, Vm58, Vi41, Am03 and Am06 (Fig. S2). Among these lineages, only Vm09 and Vm58 consistently yielded adult emergence rates above 50% (Fig. S2b,n). We therefore selected Vm09 and Vm58 for detailed genetic and functional analyses.

### Candidate mutations associated with evolution of mutualism in *E. coli* lineage Vm09

We first analyzed Vm09. Reinoculation assays of 9–11 clones from each generation showed significant improvements in adult emergence and body coloration from generation 5 onwards (Fig. S3a,b; Table S4), indicating that host-beneficial mutations rose in frequency around generation 5. We then selected four clones from each of generations 3 and 4 as pre-evolution representatives and four clones from each of generations 5, 6 and 7 as post-evolution representatives. Reinoculation confirmed the phenotypic contrast between these groups (Fig. S3c,d). Whole-genome resequencing identified 12 mutations that appeared and became predominant from generation 5, including nonsynonymous or disruptive variants in *kdpE*, *pncB*, *ydiK*, *yebK*, *yggE*, *sspA*, *waaS* and *leuP*. A nonsynonymous *rpoB* mutation arose later, in generation 6, and persisted in generation 7 (Fig. 3a,b).

**Fig. 3.**
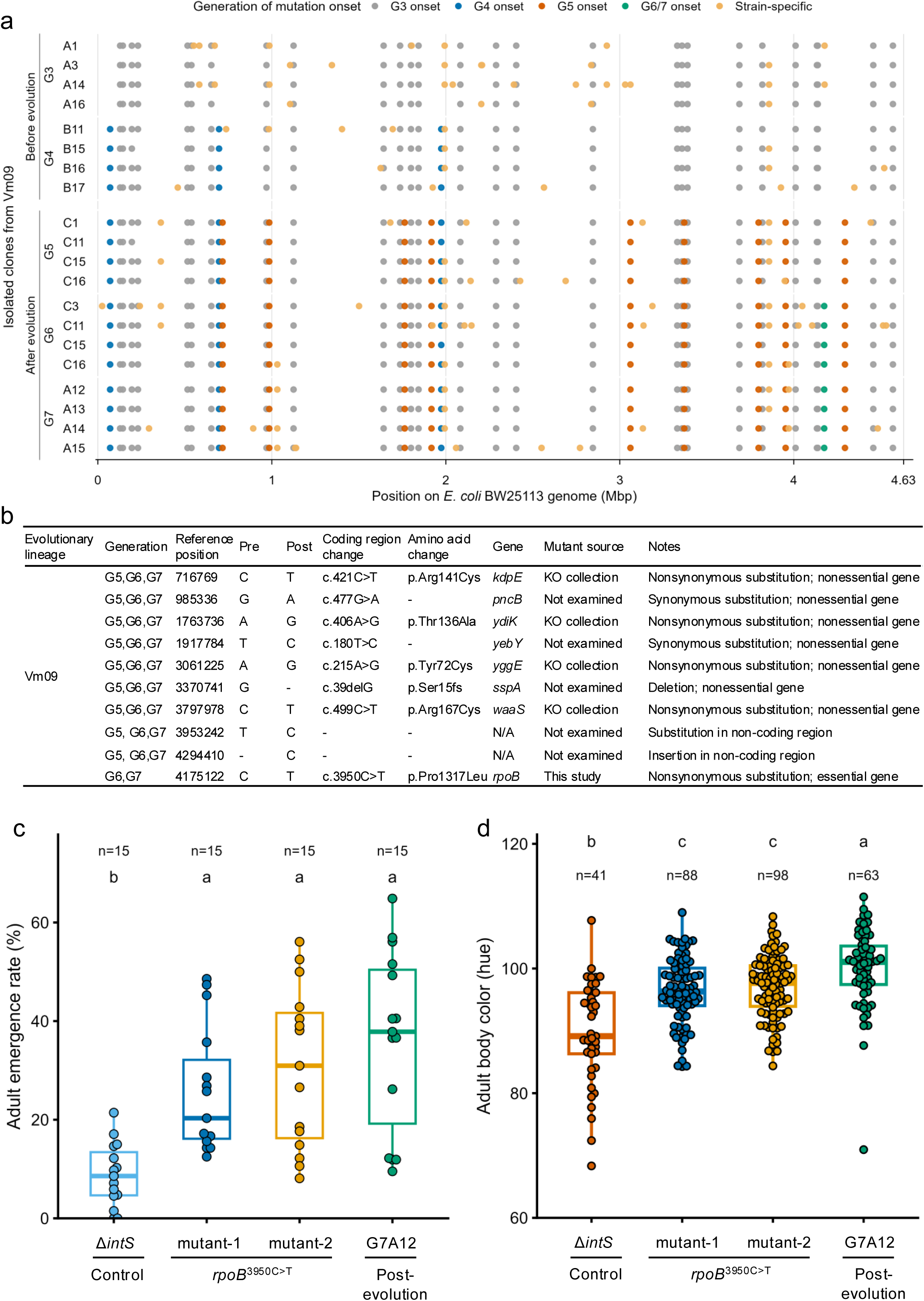
Identification and functional validation of mutations associated with host-beneficial evolution in *E. coli* lineage Vm09. (**a**) Genome-wide distribution of mutations detected in sequenced Vm09 clones. Each point marks a mutation at its position in the *E. coli* BW25113 genome; colors indicate the generation in which the mutation was first detected (G3, G4, G5, G6/7 or clone-specific). Sequenced clones and genome libraries are listed in Tables S4 and S2, respectively. (**b**) Candidate mutations associated with the evolution of host benefit in Vm09. (**c,d**) Adult emergence rate (**c**) and body coloration (**d**) of *P. stali* infected with the evolved clone G7A12 or independently generated *rpoB*^3950C>T^ mutants. Dots represent biological replicates; boxes show medians and interquartile ranges (IQRs), and whiskers extend to 1.5 × IQR. Different letters indicate significant differences (two-sided pairwise Wilcoxon rank-sum tests with Hommel correction; *P* < 0.05).

### *waaS* mutation possibly contributing to evolution of mutualism in *E. coli* lineage Vm09

Among the candidate genes identified in Vm09, Keio collection deletion mutants were available for *kdpE*, *ydiK*, *yggE* and *waaS*^23^. Reinoculation assays showed that only Δ*waaS* significantly increased adult emergence (Fig. S4), suggesting that loss of *waaS* function can improve host performance and may have contributed to the phenotypic improvement observed from generation 5. Because *waaS* participates in lipopolysaccharide core-oligosaccharide synthesis^24^, altered cell-surface structure provides one possible explanation for this effect. However, the Keio deletion does not reproduce the evolved *waaS*^499C>T^ allele; the contribution of that specific mutation therefore remains unresolved.

### The evolved *rpoB* allele confers host-beneficial phenotypes in Vm09

In Vm09, host performance improved from generation 5 and remained elevated in generations 6 and 7 (Fig. S3). The *rpoB*^3950C>T^ mutation first appeared in generation 6 (Fig. 3a,b), suggesting that it might have enhanced or stabilized the mutualistic phenotype thereafter. Because *rpoB* is essential in *E. coli* and no deletion mutant is available^23^, we reconstructed the evolved *rpoB*^3950C>T^ allele in the ancestral background. Inoculation of two independently engineered mutants significantly increased host adult emergence and improved body coloration (Fig. 3c,d). Thus, *rpoB*^3950C>T^ is sufficient to confer a substantial host-beneficial effect, although other mutations, including the earlier *waaS* allele, probably contributed to the onset of mutualism in Vm09.

### The evolved *rpoD* allele confers host-beneficial phenotypes in Vm58

We next analyzed Vm58. Reinoculation assays showed significant improvements in adult emergence and body coloration in generations 3 and 5 relative to generation 1 (Fig. S5a,b; Table S5), indicating that host-beneficial mutations arose between generations 1 and 3. Because phenotypes varied among clones, we selected eleven low-emergence clones from generation 1 and six high-emergence clones from each of generations 3 and 5 for confirmatory assays and genome sequencing (Fig. S5c,d). Comparative genomics identified seven mutations that appeared by generation 3 and persisted to generation 5, including nonsynonymous changes in *umuC*, *tsaB*, *rpoD*, *mnmE* and *rpoB* (Fig. 4a,b). We reconstructed the evolved *mnmE*^C>T^, *umuC*^1214C>T^ and *rpoD*^1712A>G^ alleles in the ancestral background, but repeated attempts to reconstruct *tsaB*^22A>G^ and *rpoB*^2137G>A^ were unsuccessful. Among the reconstructed mutants, only *rpoD*^1712A>G^ significantly increased host adult emergence (Fig. 4c,d). Thus, *rpoD*^1712A>G^ is sufficient to confer a host-beneficial effect, while possible contributions from the unreconstructed *tsaB* and *rpoB* alleles or epistatic interactions remain unresolved.

**Fig. 4.**
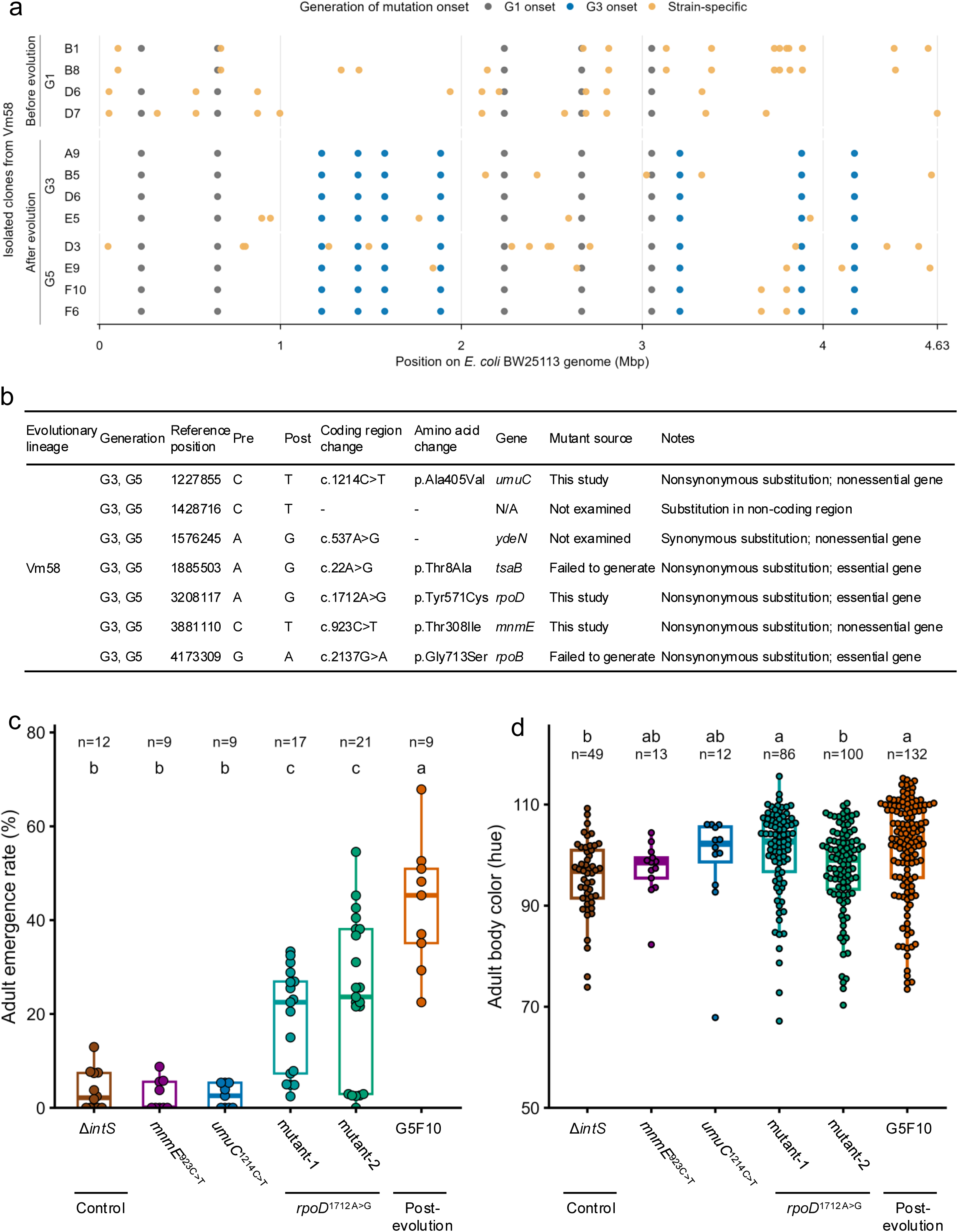
Identification and functional validation of mutations associated with host-beneficial evolution in *E. coli* lineage Vm58. (**a**) Genome-wide distribution of mutations detected in sequenced Vm58 clones. Each point marks a mutation at its position in the *E. coli* BW25113 genome; colors indicate the generation in which the mutation was first detected (G1, G3 or clone-specific). Sequenced clones and genome libraries are listed in Tables S5 and S2, respectively. (**b**) Candidate mutations associated with the evolution of host benefit in Vm58. (**c,d**) Adult emergence rate (**c**) and body coloration (**d**) of *P. stali* infected with evolved clone G5F10 or engineered *mnmE*^C>T^, *umuC*^1214C>T^ or *rpoD*^1712A>G^ mutants. Two independently generated *rpoD*^1712A>G^ mutants were tested to control for unintended background mutations. Dots represent biological replicates; boxes show medians and IQRs, and whiskers extend to 1.5 × IQR. Different letters indicate significant differences (two-sided pairwise Wilcoxon rank-sum tests with Hommel correction; *P* < 0.05).

### Transcriptomics of *E. coli* lineages Vm09 and Vm58, and *rpoB* and *rpoD* mutants

Because *rpoB* encodes the RNA polymerase β subunit^25^ and *rpoD* encodes the primary σ^70^ factor^26^, the evolved alleles were expected to alter transcription broadly. We therefore performed transcriptomic profiling of bacteria recovered from dissected symbiotic organs and compared evolved clones before and after host-beneficial phenotypes emerged with reconstructed mutant and ancestral control strains. For Vm09, the pre-evolution group comprised four clones from each of generations 3, 4 and 5 lacking *rpoB*^3950C>T^, whereas the post-evolution group comprised four generation-6 clones and seven generation-7 clones carrying the allele (Table S4). Hierarchical clustering separated the two groups, and the reconstructed *rpoB*^3950C>T^ mutants clustered with post-evolution clones (Fig. 5a). For Vm58, three generation-1 clones lacking *rpoD*^1712A>G^ were compared with four generation-5 clones carrying the allele (Table S5); the reconstructed *rpoD*^1712A>G^ mutants clustered with the latter group (Fig. 5b). Differential gene expression analysis identified hundreds of genes altered across the evolved lineages and the reconstructed mutants (Fig. S6).

**Fig. 5.**
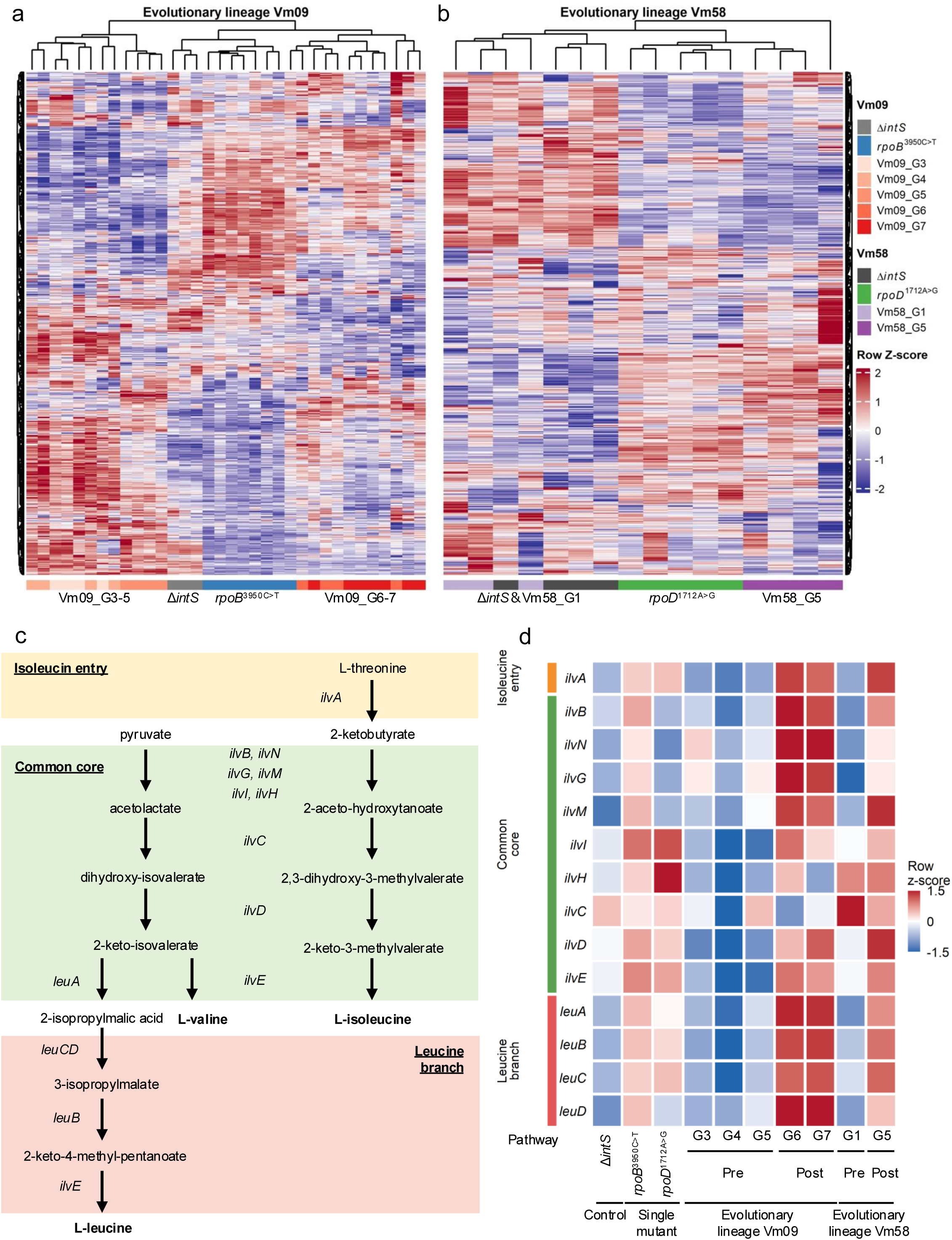
Transcriptomic changes associated with host-beneficial evolution in Vm09 and Vm58 and with reconstructed *rpoB* and *rpoD* alleles. (**a,b**) Heatmaps of the 1,000 most variable genes in Vm09 (**a**) and Vm58 (**b**), based on variance-stabilized RNA-seq counts. Expression values were standardized within each gene and are shown as row Z-scores; red and blue indicate expression above and below the gene-wise mean, respectively. Genes and samples were hierarchically clustered using Euclidean distance and complete linkage. Sample annotations indicate Δ*intS* controls, reconstructed *rpoB*^3950C>T^ mutants and Vm09 G3–G7 clones in (**a**), and Δ*intS* controls, reconstructed *rpoD*^1712A>G^ mutants and Vm58 G1 and G5 clones in (**b**). (**c**) Branched-chain amino acid biosynthesis pathways in *E. coli*, comprising the isoleucine-entry, common-core and leucine-branch pathways. (**d**) Heatmap of branched-chain amino acid biosynthesis gene expression in Vm09 before and after emergence of the *rpoB*^3950C>T^ allele, in Vm58 before and after emergence of the *rpoD*^1712A>G^ allele, and in the corresponding reconstructed mutants and Δ*intS* controls.

### Branched-chain amino acid biosynthesis genes are upregulated across independent mutualistic trajectories

Gene Ontology enrichment analysis identified functional categories associated with genes upregulated or downregulated after the evolution of mutualism and after reconstruction of the *rpoB*^3950C>T^ or *rpoD*^1712A>G^ allele (Figs. S7 and S8). In Vm58, the most markedly enriched upregulated categories were related to the biosynthesis and metabolism of branched-chain amino acids (BCAAs), valine, leucine and isoleucine (Fig. S7b). BCAA-related categories were also enriched among genes upregulated in Vm09 and in the reconstructed *rpoB*^3950C>T^ and *rpoD*^1712A>G^ mutants (Fig. S7a,c), but were not found among downregulated genes (Fig. S8). Examination of individual genes in the isoleucine-entry, common-core and leucine-branch pathways (Fig. 5c) showed broad upregulation in both Vm09 and Vm58 after mutualism evolved and in both reconstructed mutants (Fig. 5d; Fig. S9). These concordant transcriptional changes identify BCAA biosynthesis as a shared downstream response to the *rpoB* and *rpoD* evolutionary trajectories.

### Bacterial phenotypes of *E. coli* before and after evolution of mutualism and with and without beneficial *rpoB* and *rpoD* mutations

We examined whether mutualism evolution and reconstruction of the *rpoB*^3950C>T^ or *rpoD*^1712A>G^ allele were accompanied by changes in bacterial growth, morphology and motility. Most evolved clones and reconstructed mutants grew more slowly in liquid LB medium than the control strains (Fig. S10a) and exhibited smaller cells (Fig. S10b–d). Swimming motility was also strongly reduced in a subset of evolved clones and in the reconstructed *rpoB* mutants (Fig. S10e–g). These changes resemble the reduced cell size and motility previously observed in mutualistic *E. coli* evolved with *P. stali* due to CCR mutations^14^. Together, the results show that host-beneficial evolution was repeatedly associated with attenuation of traits characteristic of growth outside the host. Meanwhile, whether these changes directly improve within-host performance remains to be tested.

## Discussion

In this study, we established 144 experimental evolutionary *E. coli* lineages in association with the stinkbug *P. stali* and identified six lineages with reproducible host-beneficial phenotypes. All six originated from the 73 hypermutator Δ*mutS* lineages, whereas none arose among the 71 control Δ*intS* lineages, consistent with an increased supply of adaptive mutations in the hypermutator background. Host-beneficial phenotypes emerged within one to five host generations. Four of the six lineages independently acquired *cyaA* mutations, and three tested lineages reproducibly improved host performance. Notably, CCR-associated trajectories occurred in four of 73 Δ*mutS* lineages in this study and two of 19 in our previous study^14^. These repeated outcomes identify CCR disruption as an accessible and recurrent route to *P. stali*–*E. coli* mutualism under the experimental conditions adopted here.

In addition, we identified two evolved *E. coli* lineages that improved host performance without CCR mutations. Reconstruction experiments showed that the evolved *rpoB*^3950C>T^ and *rpoD*^1712A>G^ alleles were each sufficient to confer host-beneficial effects in the ancestral genetic background. These alleles affect core components of the transcriptional machinery: *rpoB* encodes the β subunit of RNA polymerase, whereas *rpoD* encodes the primary σ^70^ factor. Together with the recurrent *cyaA* mutations, these findings reveal convergence at a higher functional level on global transcriptional regulation. Meanwhile, neither allele alone necessarily explains the full evolutionary history of its lineage: *rpoB*^3950C>T^ arose after the initial improvement in Vm09, and unreconstructed mutations or epistatic interactions may also have contributed in Vm58. Taken together, global transcriptional regulators represent recurrent, large-effect targets of mutualistic evolution under the conditions examined here.

Mutations in global regulatory genes frequently arise during microbial experimental evolution under diverse selective pressures, including temperature stress, antibiotic exposure and nutrient limitation^27–35^. The recurrent recovery of *cyaA*, *rpoB* and *rpoD* mutations in the *P. stali*–*E. coli* system is consistent with this broader pattern. Such mutations may be favored because a single genetic change can alter many traits simultaneously, allowing rapid adaptation to a novel within-host environment. This interpretation remains a hypothesis, however, because the present experiments did not directly compare the selective advantages, mutational target sizes or competitive dynamics of global-regulator and downstream-effector alleles.

How can global regulatory mutations make *E. coli* beneficial to *P. stali*? In the case of CCR disruption, a previous work identified reduced *tnaA* expression as a downstream mechanism that contributes to host fitness via increased tryptophan availability and suppressed indole production^15^. Here, Vm09 and Vm58, together with reconstructed *rpoB*^3950C>T^ and *rpoD*^1712A>G^ mutants, consistently showed increased expression of genes involved in biosynthesis of essential BCAAs, namely valine, leucine and isoleucine. On the ground that plant-sucking hemipteran insects such as aphids, cicadas, leafhoppers and stinkbugs commonly depend on symbionts for supply of essential amino acids that are scarce in plant sap^36–40^, enhanced BCAA biosynthetic capacity provides a plausible explanation for the observed host benefits. This interpretation is consistent with the beneficial effects of *E. coli* mutations affecting tryptophan and methionine metabolism^15,16^. Nevertheless, transcript abundance does not guarantee increased BCAA production or transfer to the host. Direct metabolite measurements and targeted perturbation of BCAA biosynthesis genes will provide further insight into whether amino acid provisioning mediates the effects of *rpoB* and *rpoD*.

Based on the present and previous experiments with the *P. stali*–*E. coli* system^13–16^, we propose a model for the early stages of mutualism under stable laboratory conditions (Fig. 6). A bacterium that initially provides little benefit may acquire regulatory mutations that increase the production or availability of host-limiting metabolites. Because global regulators affect many traits, the same mutations may also reduce functions associated with growth outside the host. Subsequent selection could stabilize host-beneficial phenotypes and permit the gradual erosion of functions that are dispensable within the host, paralleling patterns of genome reduction in long-term insect symbionts^6–9^. This model links rapid regulatory adaptation to later host dependence, but the experiments reported here capture only the earliest transition to measurable host benefit. Processes expected at later evolutionary stages, such as genetic accommodation, genome degradation and the evolution of obligate dependence, could be explored in future long-term experimental evolution studies.

**Fig. 6.**
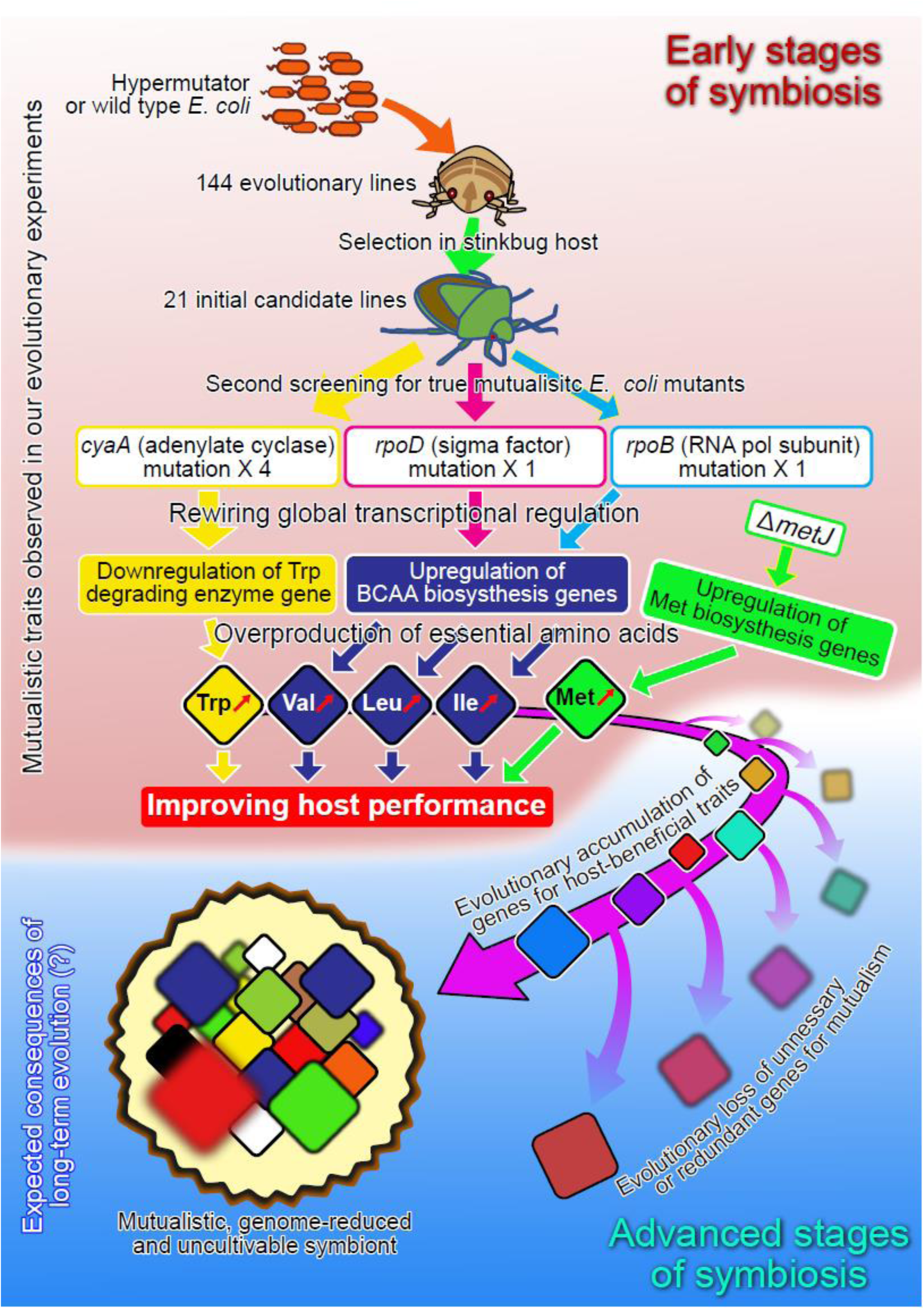
Proposed model for the early evolution of *P. stali*–*E. coli* mutualism through global transcriptional rewiring. Mutations in global regulator**y** genes alter downstream pathways that enhance host-beneficial functions while attenuating traits associated with growth outside the host.

Why were global regulators repeatedly recovered whereas known downstream effectors such as *tnaA* were not? One possibility is that pleiotropic regulatory mutations generate composite phenotypes favored inside the host: they may increase nutritional value while reducing costly free-living traits or improving persistence within the symbiotic organ^11,12^. Consistent with this interpretation, *cyaA*, *rpoB* and *rpoD* mutants showed reduced growth, smaller cell size or impaired motility^14^ (see Fig. S10), whereas the *tnaA* mutant did not show such phenotypes^15^ (see Fig. S11). However, alternative explanations include differences in mutational target size, allele-specific fitness effects, clonal interference and the screening criteria used to identify beneficial lineages. Distinguishing among these mechanisms will require direct competition experiments inside and outside the host and systematic comparisons of engineered regulator and effector alleles.

The *P. stali*–*E. coli* evolutionary trajectory toward mutualism is consistent with the proposal that early symbiotic evolution can be driven by mutations with large pleiotropic effects^12,41,42^. Global regulatory mutations have also arisen during experimental or early-stage host associations such as the squid–*Aliivibrio* luminescent symbiosis^43^, a nematode– *Pseudomonas* association^44^, the experimental conversion of a plant pathogen into a legume symbiont^45^, and amoeba-associated *Parachlamydia* passaging^46^. Together, these systems suggest that regulatory-network reprogramming is a recurrent route to rapid host adaptation. Whether the same mutations subsequently promote obligate dependence, however, remains unresolved and is likely to depend on transmission mode, ecological stability and the fitness costs of pleiotropy outside the host.

These findings derive from experimental evolution under controlled laboratory conditions and should therefore be extrapolated to the origin of mutualism in nature with caution. Notably, disruptive mutations in *cyaA* and *crp* have generally not been detected in natural stinkbug symbionts at early stages of symbiotic evolution^15^. Global regulatory genes are important for adaptation to environmental variability; mutations in these genes may have been favored under the constant laboratory conditions used here, whereas they may be disadvantageous in the more variable environments encountered in nature. Thus, whether comparable regulatory changes contribute to the early evolution of natural insect–bacterium mutualisms remains to be verified in future studies.

In conclusion, large-scale experimental evolution revealed that independent routes to *P. stali*–*E. coli* mutualism repeatedly converge on global transcriptional regulation. Recurrent *cyaA* mutations and host-beneficial *rpoB* and *rpoD* alleles demonstrate that regulatory-network reprogramming can rapidly generate beneficial bacterial phenotypes, while the shared upregulation of essetial amino acid biosynthesis genes provides a plausible mechanistic link to host nutrition. These findings identify global transcriptional regulators as accessible early targets in the experimental evolution of mutualism and suggest how ordinary bacteria can rapidly acquire functions that support the nutritionally specialized, plant-sucking lifestyle of host insects. Direct metabolic measurements and tests under ecologically variable conditions will now be essential to determine how often comparable transitions occur in nature.

## Methods

### Insect material, bacterial strains, evolved isolates and sequencing libraries

An inbred laboratory strain of the stinkbug *P. stali* was used throughout this study^47^. The bacterial strains, plasmids and oligonucleotides are listed in Table S7. Evolved *E. coli* isolates from lineages Vm09 and Vm58 used for reinoculation assays, genomic analyses and transcriptomic analyses are listed in Tables S4 and S5. DNA-and RNA-sequencing libraries are summarized in Tables S2 and S6, respectively.

### Insect rearing, bacterial inoculation and host-performance analyses

Insects were reared, rendered symbiont-free and inoculated with bacteria as described previously^15^. Cohorts were maintained for six weeks after egg collection, at which point adult emergence and body coloration were recorded. Adults were anaesthetized by overnight refrigeration and scanned dorsally using a GT-X980 flatbed scanner (Epson). Body coloration was quantified from the resulting images using Natsumushi v1.10^48^.

### Experimental evolution of *P. stali*–*E. coli* association

Large-scale evolution experiments were conducted under vertical-transmission and artificial-inoculation regimes (Fig. 1). Independent lineages were founded by separate inoculation events using Δ*mutS* or Δ*intS E. coli* and approximately 84 newly hatched nymphs per lineage. The first inoculated cohort was designated generation 1. The vertical-transmission experiment comprised 58 Δ*mutS* and 56 Δ*intS* lineages, whereas the artificial-inoculation experiment comprised 15 lineages of each genotype. Initial sterilization and inoculation followed previously published procedures^14^. In each generation, insects were reared to adulthood and up to ten mating pairs were established 7–10 days after adult emergence. Egg masses produced over the subsequent 2–3 weeks were assigned unique maternal identifiers, and the five females with the highest egg production were retained. Thus, selection acted on bacterial effects on host survival, coloration, reproductive output and lineage persistence, while repeated passage through individual females imposed population bottlenecks. Some evolutionary lineages produced few or no adults and became extinct, which could be restarted from cryopreserved symbiotic organ homogenates. After oviposition, the symbiotic organ from each selected female was dissected and homogenized individually in 400 μl phosphate-buffered saline. Most of each homogenate was mixed with glycerol and stored at −80 °C; the remainder was used for contamination screening. Universal bacterial 16S rRNA and *E. coli*-specific Tn5 copy numbers were quantified by qPCR, and diluted homogenates were plated on LB agar with and without kanamycin. Only offspring from females judged free of detectable contamination were retained for the next generation, following the quality-control procedures described previously^14^.

### Targeted sequencing of CCR-associated genes

Candidate *E. coli* lineages were screened for mutations in *cyaA* and *crp*, two key components of the CCR pathway. The target loci were amplified using Tks Gflex DNA Polymerase (Takara Bio) and the primers listed in Table S7. The resulting PCR products were subjected to Sanger sequencing. Sequence reads were aligned to the corresponding loci on the genome of ancestral *E. coli* strain to identify nucleotide substitutions and insertion–deletion variants.

### Genome sequencing and mutation detection

For the *cyaA*-mutant lineages Vm07, Vm18 and Vm57, genomic DNA was prepared from cryopreserved samples collected at successive generations (Tables S2 and S3). For Vm09 and Vm58, genomic DNA was prepared from representative evolved clones isolated at each generation (Tables S2, S4 and S5). DNA was purified using the QuickGene-AutoS DNA Blood Kit (Kurabo Industries) and sequenced by Azenta Life Sciences (GENEWIZ) on an Illumina NovaSeq platform in paired-end 150-bp mode, with a target yield of 2 Gb per sample. After quality filtering, reads were mapped to the *E. coli* BW25113 reference genome (CP009273.1) using CLC Genomics Workbench v25.0.0. Because the experiment was initiated with a laboratory-maintained BW25113 Δ*mutS* strain that could differ from the public reference sequence, five ancestral isolates (G01F1–G01F5) were also sequenced (Table S2). Variants shared by all five ancestral isolates were classified as pre-existing background variants. De novo mutations were identified as variants absent from the ancestral background that emerged during passage within each lineage. Data processing and visualization were performed in R v4.4.0 and RStudio 2026.01.2 Build 418.

### Construction of *E. coli* mutants

The evolved *rpoB*^3950C>T^ and *rpoD*^1712A>G^ alleles were introduced into *E. coli* BW25113 by multiplex automated genome engineering method using 90-mer DNA oligonucleotides whose first five nucleotides were phosphorothioated^49^ (Table S7). Candidate clones were screened by PCR, and the introduced alleles were verified by sequencing. Two independently generated mutants were obtained for each allele and tested to control for unintended background mutations. An *rpsL* K42T counter-selection background was constructed as described previously^50^. The *mnmE*^C>T^ and *umuC*^1214C>T^ alleles were introduced into this background by λ-Red homologous recombination using pRed/ET (Gene Bridges) as described previously^51^. All mutagenic and screening oligonucleotides are listed in Table S7. Despite repeated attempts with both approaches, *rpoB*^2137G>A^ and *tsaB*^22A>G^ could not be recovered.

### Transcriptomic analyses

Symbiotic organs were dissected from *E. coli*-infected adults 5–7 days after adult emergence and used for RNA extraction. Libraries were prepared with the TruSeq Stranded mRNA Sample Preparation Kit and sequenced by Azenta Life Sciences (GENEWIZ) on an Illumina NovaSeq platform in paired-end 150-bp mode. Base calling and demultiplexing followed the provider’s standard pipeline. After quality filtering, reads were mapped to the *E. coli* BW25113 reference genome (NZ_CP009273) and counted using CLC Genomics Workbench v25.0.0. Differential expression was analysed with DESeq2; genes with an adjusted *P* value < 0.05 and an absolute log_2_ fold change of at least 0.5 were classified as differentially expressed. Variance-stabilized counts were used for hierarchical clustering and heatmaps. Gene Ontology Biological Process enrichment was tested separately for upregulated and downregulated genes, with correction for multiple testing.

### Measurement of bacterial growth, morphology and motility

Glycerol stocks were inoculated into 3 ml LB medium containing 5 g l^−1^ yeast extract, 10 g l^−1^ tryptone and 5 g l^−1^ NaCl and cultured overnight at 25 °C with shaking. For growth measurements, cultures were diluted to an initial OD_600_ of 0.001 in 200 μl LB medium and incubated in an Infinite 200 Pro F Plex microplate reader (Tecan); absorbance at 590 nm was recorded every 10 min with intermittent shaking. For cell-size and motility measurements, overnight cultures were diluted into fresh LB medium and grown at 25 °C to OD_600_ 0.9–1.2. Cells were imaged by phase-contrast microscopy using an IX83 microscope (Olympus) for morphology and by dark-field microscopy using a CX22 microscope (Olympus) at 23 °C for motility. Images and one-second videos were acquired with a DMK33UX178 CMOS camera (The Imaging Source).

Cell dimensions were quantified in ImageJ v1.54r, and trajectories were analyzed with the TrackMate plugin. Numbers of independent cultures, fields and cells analyzed, together with statistical procedures, are reported in the corresponding figure legends and source data.

## Data availability

All DNA and RNA sequencing data produced in this study were deposited in the DNA Data Bank of Japan (DDBJ) Sequence Read Archive (Tables S2 and S6). The data have been deposited with links to BioProject accession no. PRJDB42872 in the DDBJ BioProject database. Source data are provided with this paper.

## Acknowledgments

We thank Yoshiko Ishii, Naomi Shimizu, Tomoko Tanaka, Machiko Oikawa, Tomoko Matsushita, Noriko Yamane, Tetsuhiro Hachikawa, Kenyu Kiyomiya, Yumi Kanazawa, Sakiko Toyoda, Mariko Taguchi, Mizuko Ozaki, Masae Takashima and Maria Murakami for insect rearing and screening. This study was supported by the Japan Science and Technology Agency ERATO grant number JPMJER1902 (T.F. and R.K.) and CREST grant number JPMJCR25B4 (T.F.), and the Japan Society for the Promotion of Science (JSPS) KAKENHI grant number JP24H02298 (T.F. and M.M.).

## Author contributions

R.K. and T.F. conceived the project and designed the experiments. Y.W. and R.S. conducted most of the experimental works including bacterial infection, insect rearing, bacterial mutant generation and screening, host fitness evaluation, and others. Y.W., Y.N., R.K. and T.N. performed large scale evolutionary experiments of *P. stali*–*E. coli* symbiosis. Y.W., R.K., R.S. and M.Mo. conducted transcriptomic analyses, whereas Y.W., R.S. and R.K. performed genomic analyses. M.Mi. measured bacterial phenotypes. Y.W. and T.F. wrote the paper with input from all authors. All authors approved the final version of the manuscript.

## SUPPLEMENTARY FIGURE CAPTIONS

**Fig. S1.**
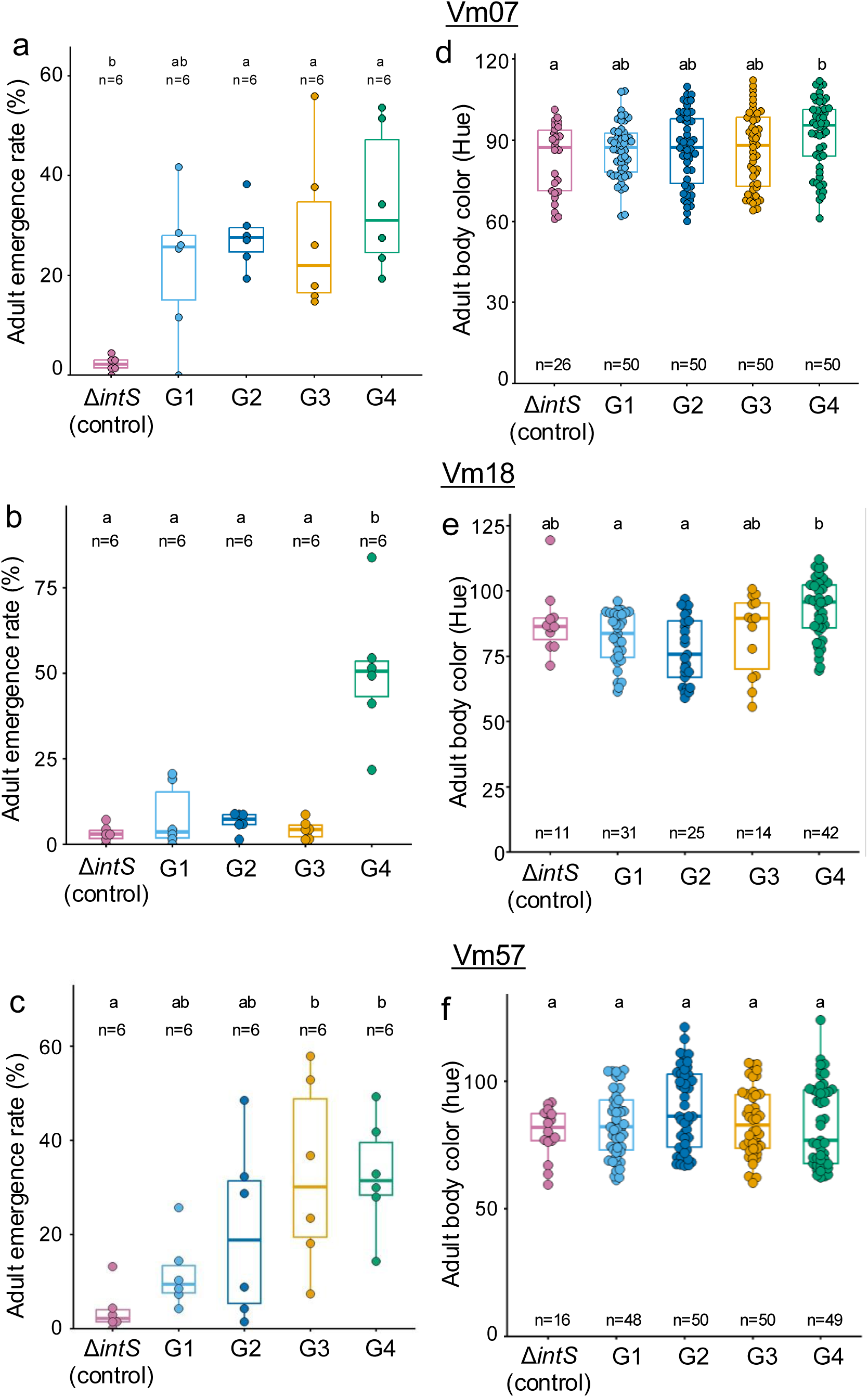
Host phenotypes following infection with evolved *E. coli* lineages carrying independent *cyaA* mutations. (**a–c**) Adult emergence rates. (**d–f**) Adult body coloration. (**a,d**) Vm07, carrying *cyaA*^512dupG^. (**b,e**) Vm18, carrying *cyaA*^1840C>T^. (**c,f**) Vm57, carrying *cyaA*^2131T>C^. Dots represent biological replicates; boxes show medians and IQRs, and whiskers extend to 1.5 × IQR. Different letters indicate significant differences (two-sided pairwise Wilcoxon rank-sum tests with Hommel correction; *P* < 0.05).

**Fig. S2.**
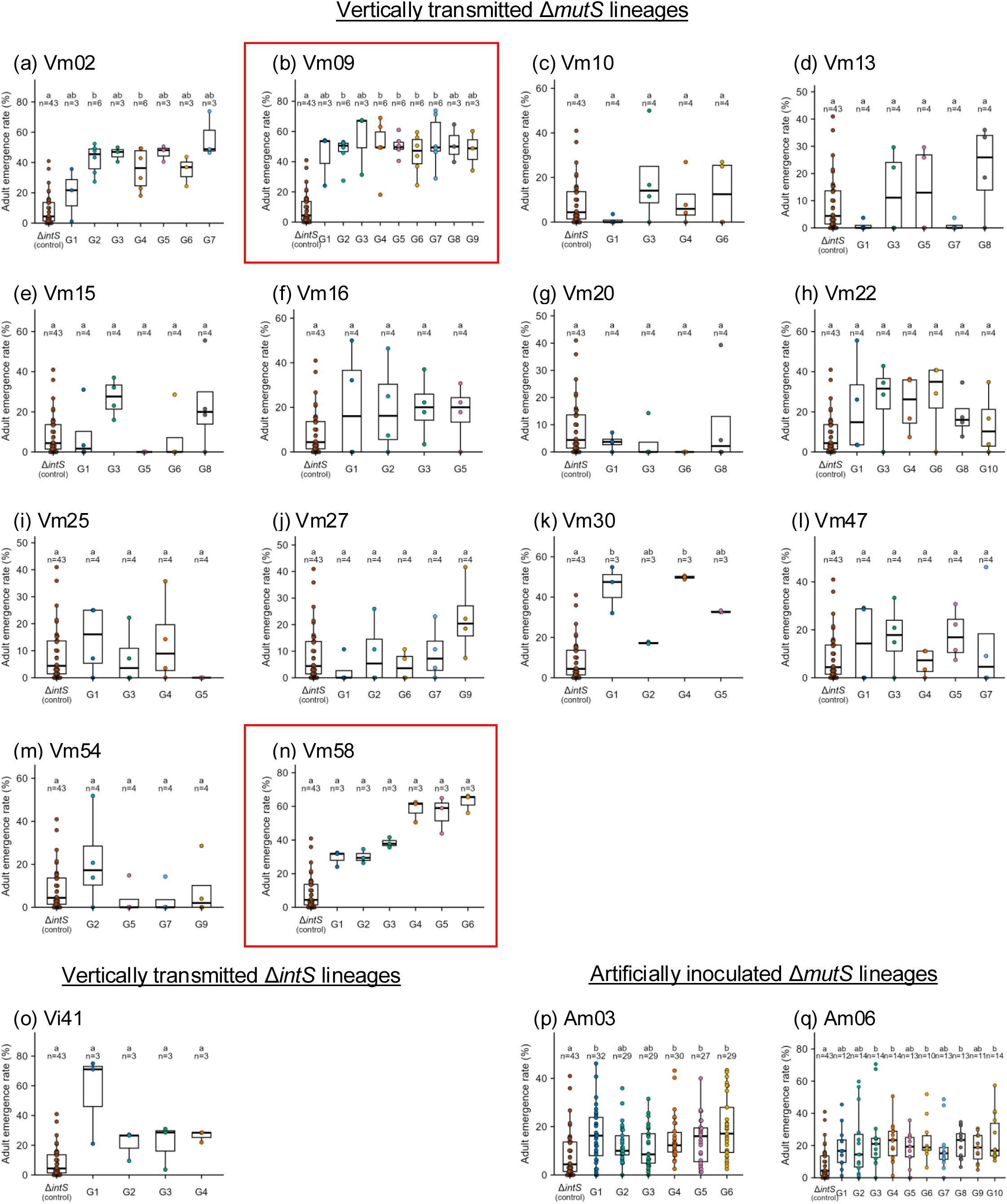
Adult emergence of *P. stali* infected with candidate evolved *E. coli* lineages. (**a**) Vm02; (**b**) Vm09; (**c**) Vm10; (**d**) Vm13; (**e**) Vm15; (**f**) Vm16; (**g**) Vm20; (**h**) Vm22; (**i**) Vm25; (**j**) Vm27; (**k**) Vm30; (**l**) Vm47; (**m**) Vm54; (**n**) Vm58; (**o**) Vi41; (**p**) Am03; (**q**) Am06. Red boxes highlight Vm09 and Vm58, which were selected for detailed analysis. Dots represent biological replicates; boxes show medians and IQRs, and whiskers extend to 1.5 × IQR. Different letters indicate significant differences (two-sided pairwise Wilcoxon rank-sum tests with Hommel correction; *P* < 0.05).

**Fig. S3.**
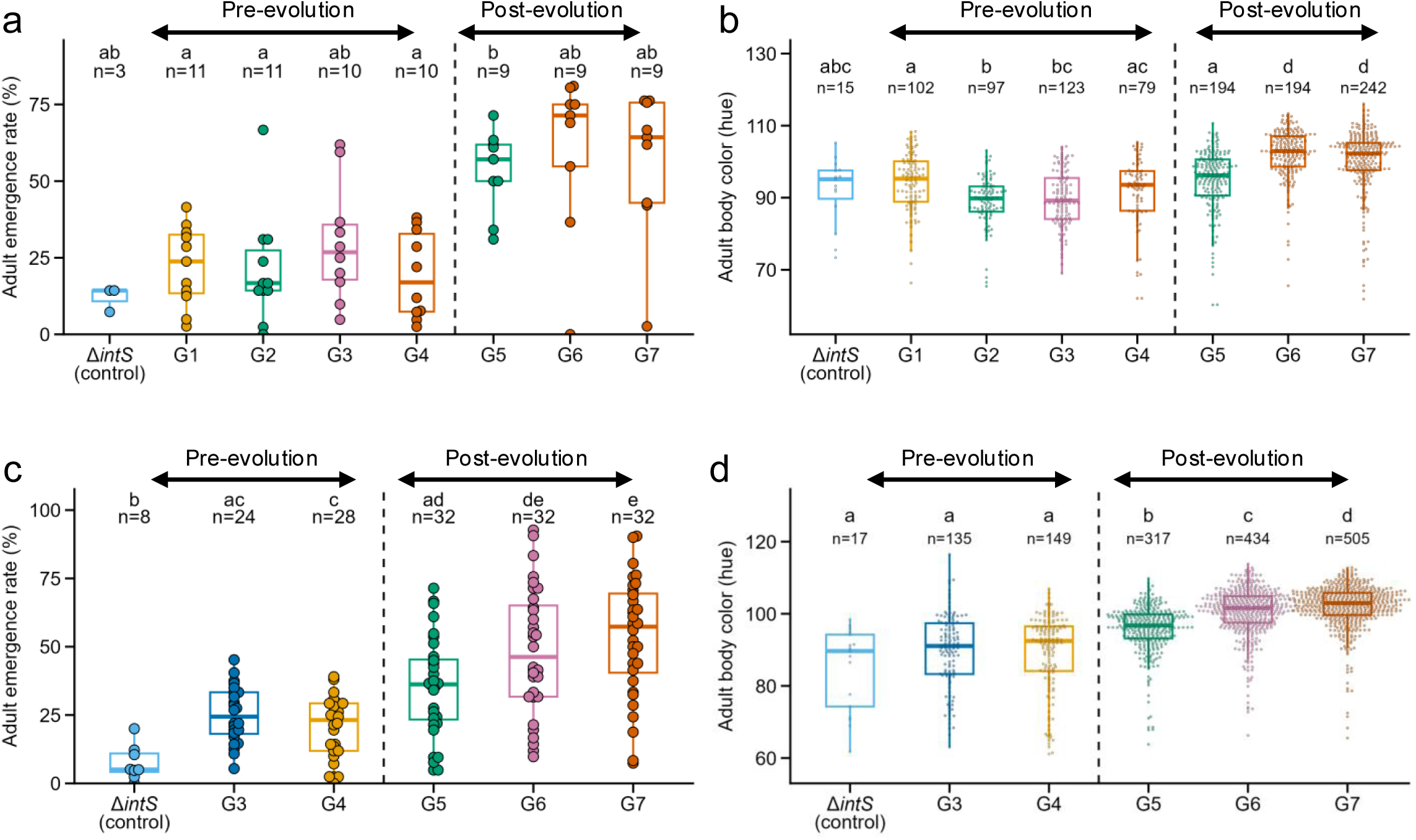
Host phenotypes following infection with *E. coli* clones isolated from Vm09. (**a,b**) Initial screen of adult emergence (**a**) and body coloration (**b**) in insects infected with clones from successive Vm09 generations, showing that host-beneficial clones became prevalent from G5. (**c,d**) Confirmatory analysis of adult emergence (**c**) and body coloration (**d**) using selected pre-evolution clones from G3 and G4 and post-evolution clones from G5– G7. Four clones with the lowest adult emergence rates were selected from each of G3 and G4, and four clones with the highest rates were selected from each of G5–G7 (Table S4). Dots represent biological replicates; boxes show medians and IQRs, and whiskers extend to 1.5 × IQR. Different letters indicate significant differences (two-sided pairwise Wilcoxon rank-sum tests with Hommel correction; *P* < 0.05).

**Fig. S4.**
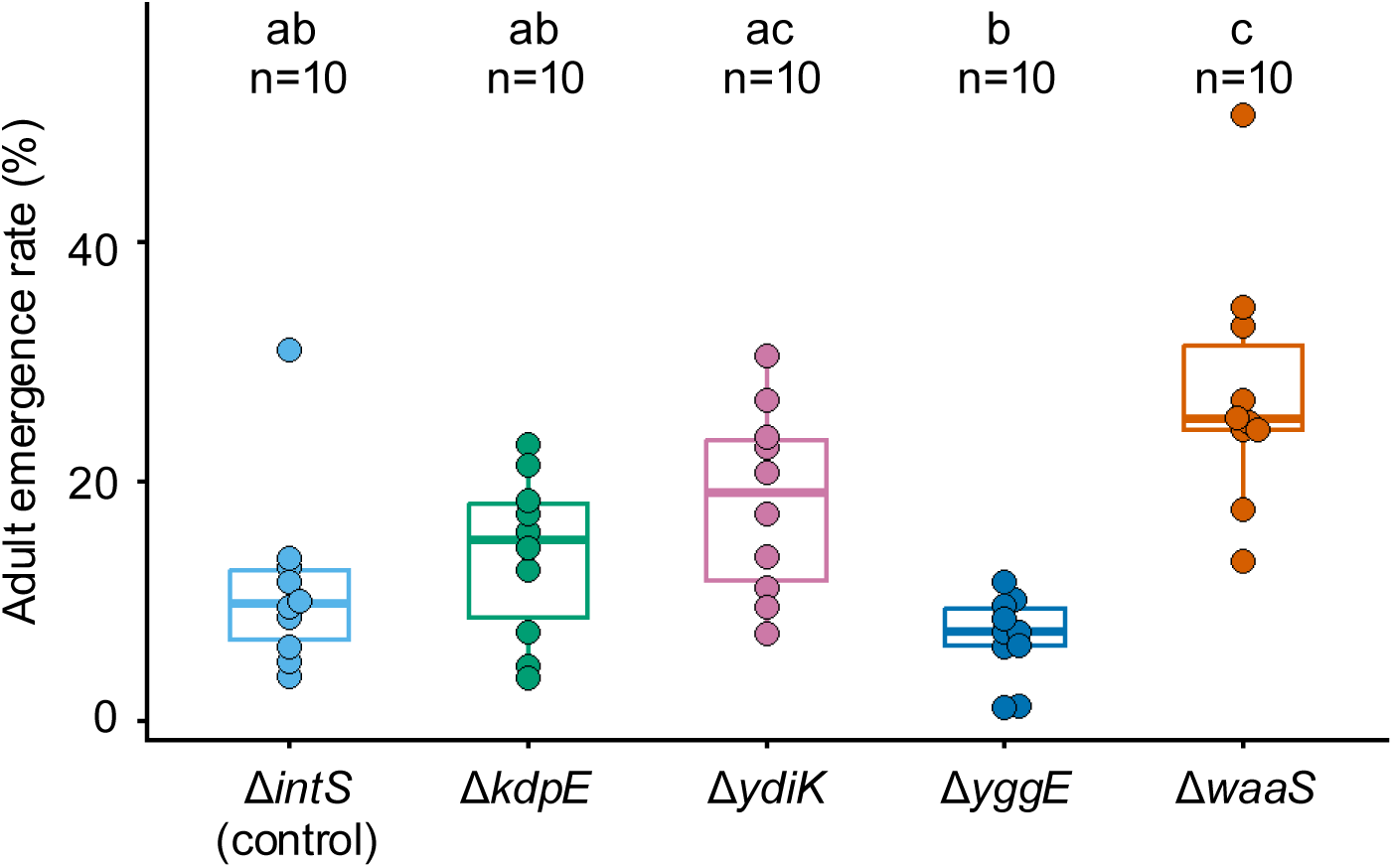
Functional testing of candidate genes associated with host-beneficial evolution in Vm09. Keio collection mutants Δ*kdpE*, Δ*ydiK*, Δ*yggE* and Δ*waaS*, corresponding to candidate genes in Fig. 3b, were inoculated to symbiont-free *P. stali* nymphs. Dots represent biological replicates; boxes show medians and IQRs, and whiskers extend to 1.5 × IQR. Different letters indicate significant differences (two-sided pairwise Wilcoxon rank-sum tests with Hommel correction; *P* < 0.05).

**Fig. S5.**
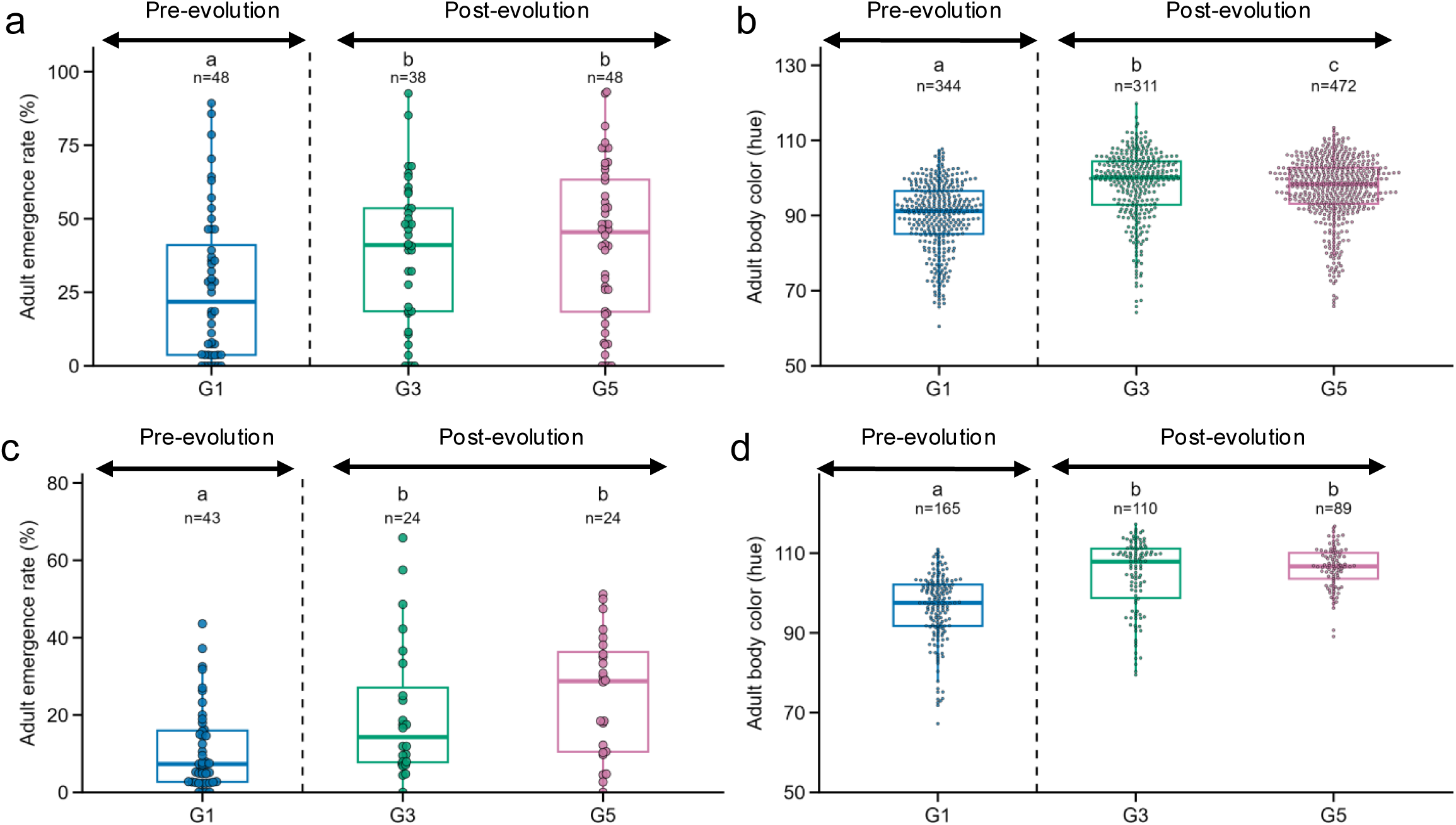
Host phenotypes following infection with *E. coli* clones isolated from Vm58. (**a,b**) Initial screen of adult emergence (**a**) and body coloration (**b**) in insects infected with clones from successive Vm58 generations, indicating that host-beneficial clones became prevalent by G3. (**c,d**) Confirmatory analysis of adult emergence (**c**) and body coloration (**d**) using selected pre-evolution clones from G1 and post-evolution clones from G3 and G5. Six clones with the lowest adult emergence rates were selected from G1, and six clones with the highest rates were selected from each of G3 and G5 (Table S5). Dots represent biological replicates; boxes show medians and IQRs, and whiskers extend to 1.5 × IQR. Different letters indicate significant differences (two-sided pairwise Wilcoxon rank-sum tests with Hommel correction; *P* < 0.05).

**Fig. S6.**
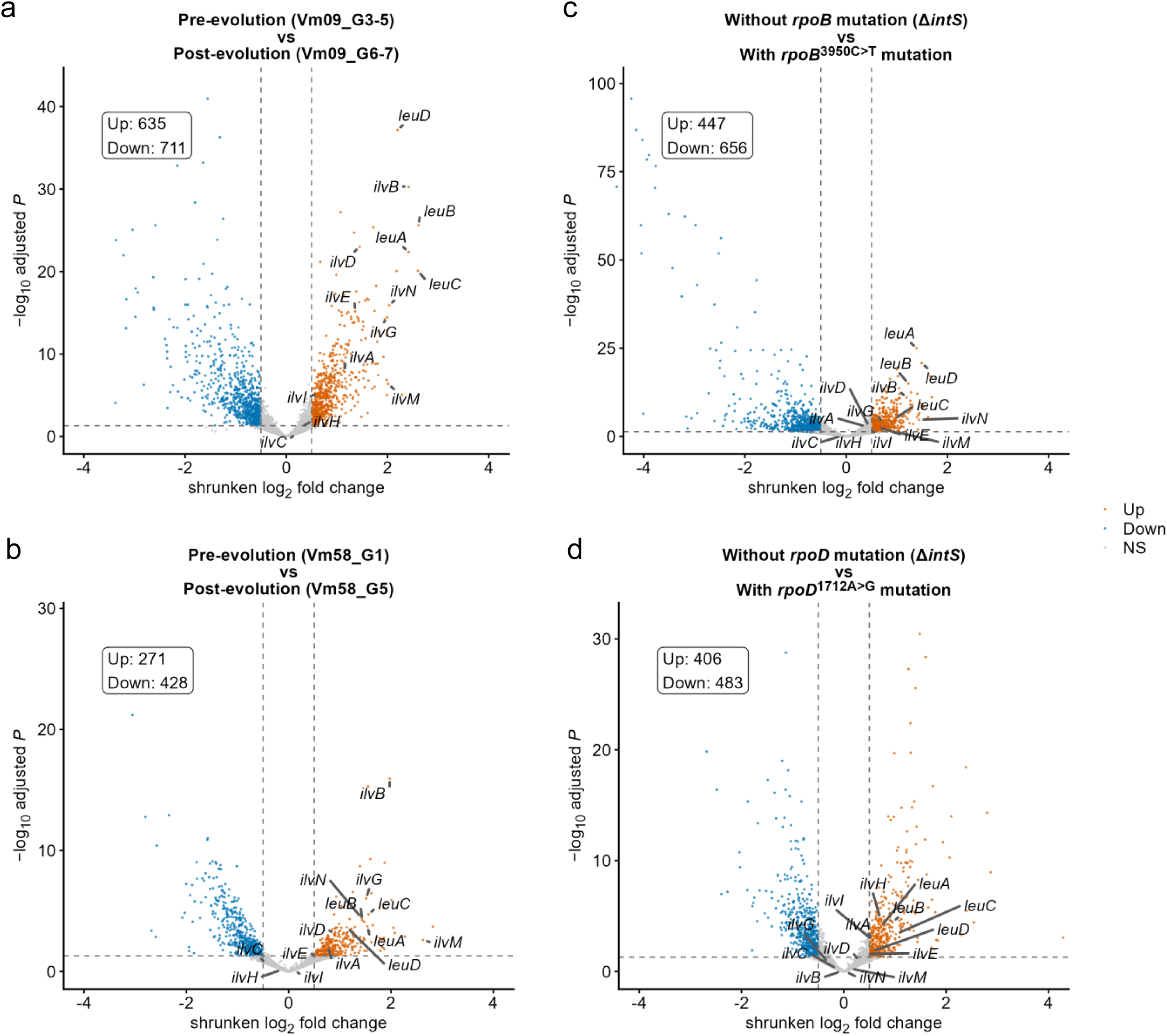
Differential gene expression associated with host-beneficial evolution and reconstructed *rpoB* and *rpoD* alleles. Volcano plots compare (**a**) Vm09 G3–G5 versus G6– G7, (**b**) Vm58 G1 versus G5, (**c**) the *rpoB*^3950C>T^ mutant versus its control and (**d**) the *rpoD*^1712A>G^ mutant versus its control. Each point represents one gene; the x axis shows log_2_ fold change and the y axis shows −log_10_-transformed adjusted *P* values. Dashed lines indicate |log_2_ fold change| ≥ 0.5 and adjusted *P* < 0.05. Numbers of upregulated and downregulated genes are shown in each panel; genes involved in branched-chain amino acid biosynthesis are highlighted.

**Fig. S7.**
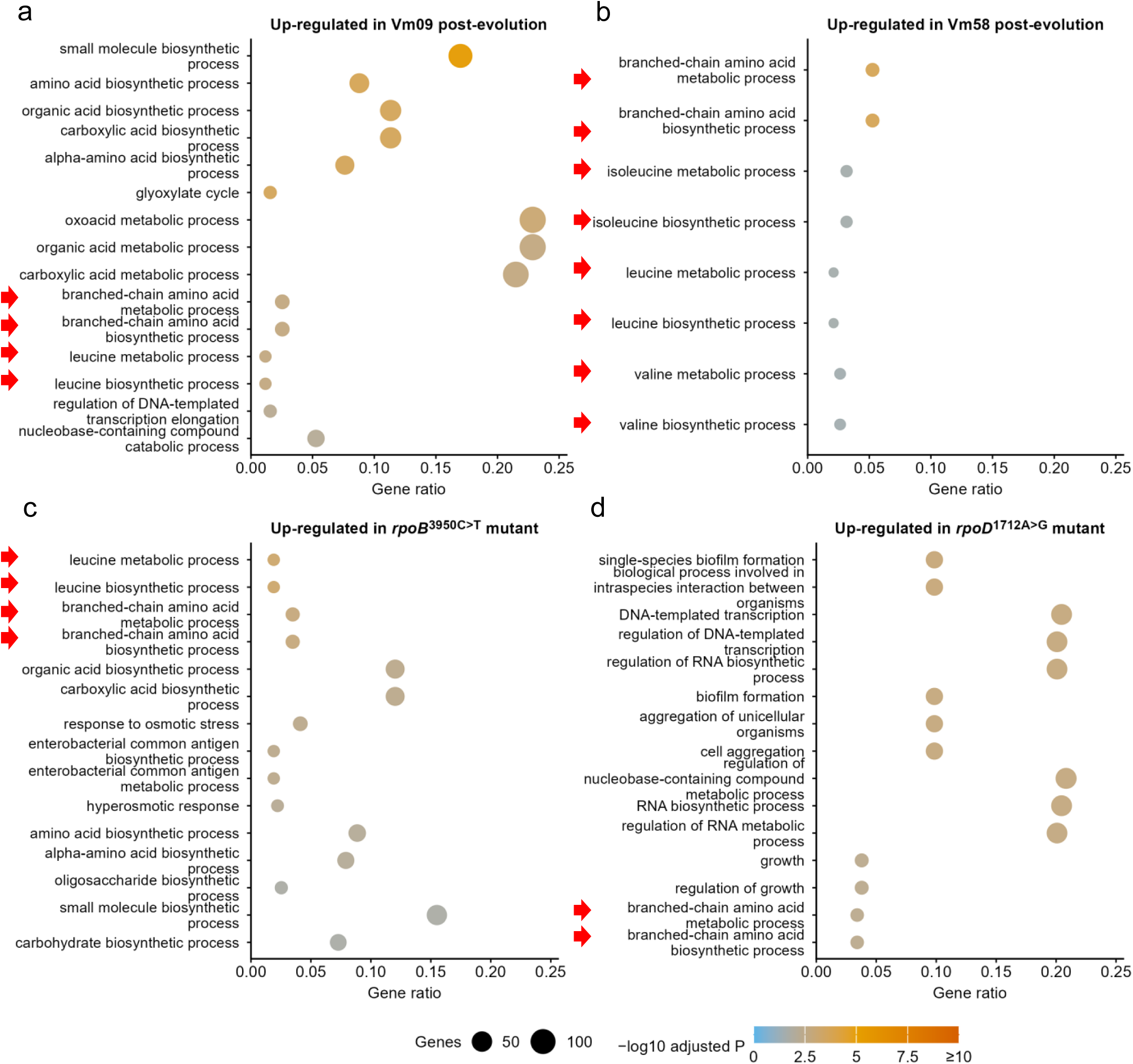
Gene Ontology enrichment among upregulated genes in Vm09, Vm58 and reconstructed *rpoB* and *rpoD* mutants. (**a**) Vm09 after host-beneficial evolution; (**b**) Vm58 after host-beneficial evolution; (**c**) reconstructed *rpoB*^3950C>T^ mutant; (**d**) reconstructed *rpoD*^1712A>G^ mutant. Enrichment was calculated from DESeq2-defined differentially expressed genes. Dot size denotes the number of genes assigned to each Gene Ontology Biological Process term, and dot color denotes the adjusted *P* value. Red arrows indicate terms related to branched-chain amino acid metabolism.

**Fig. S8.**
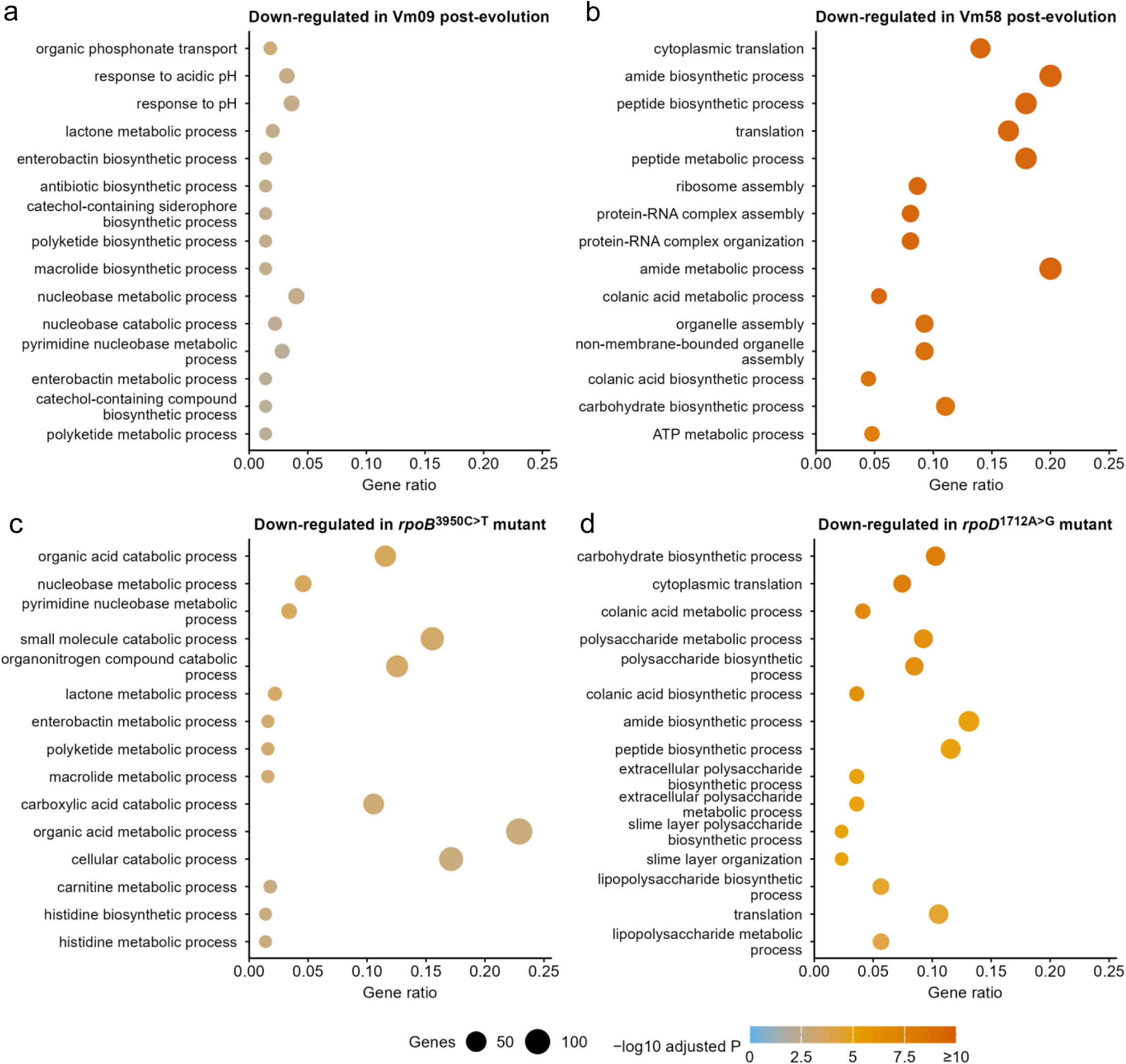
Gene Ontology enrichment among downregulated genes in Vm09, Vm58 and reconstructed *rpoB* and *rpoD* mutants. (**a**) Vm09 after host-beneficial evolution; (**b**) Vm58 after host-beneficial evolution; (**c**) reconstructed *rpoB*^3950C>T^ mutant; (**d**) reconstructed *rpoD*^1712A>G^ mutant. Enrichment was calculated from DESeq2-defined differentially expressed genes. Dot size denotes the number of genes assigned to each Gene Ontology Biological Process term, and dot color denotes the adjusted *P* value.

**Fig. S9.**
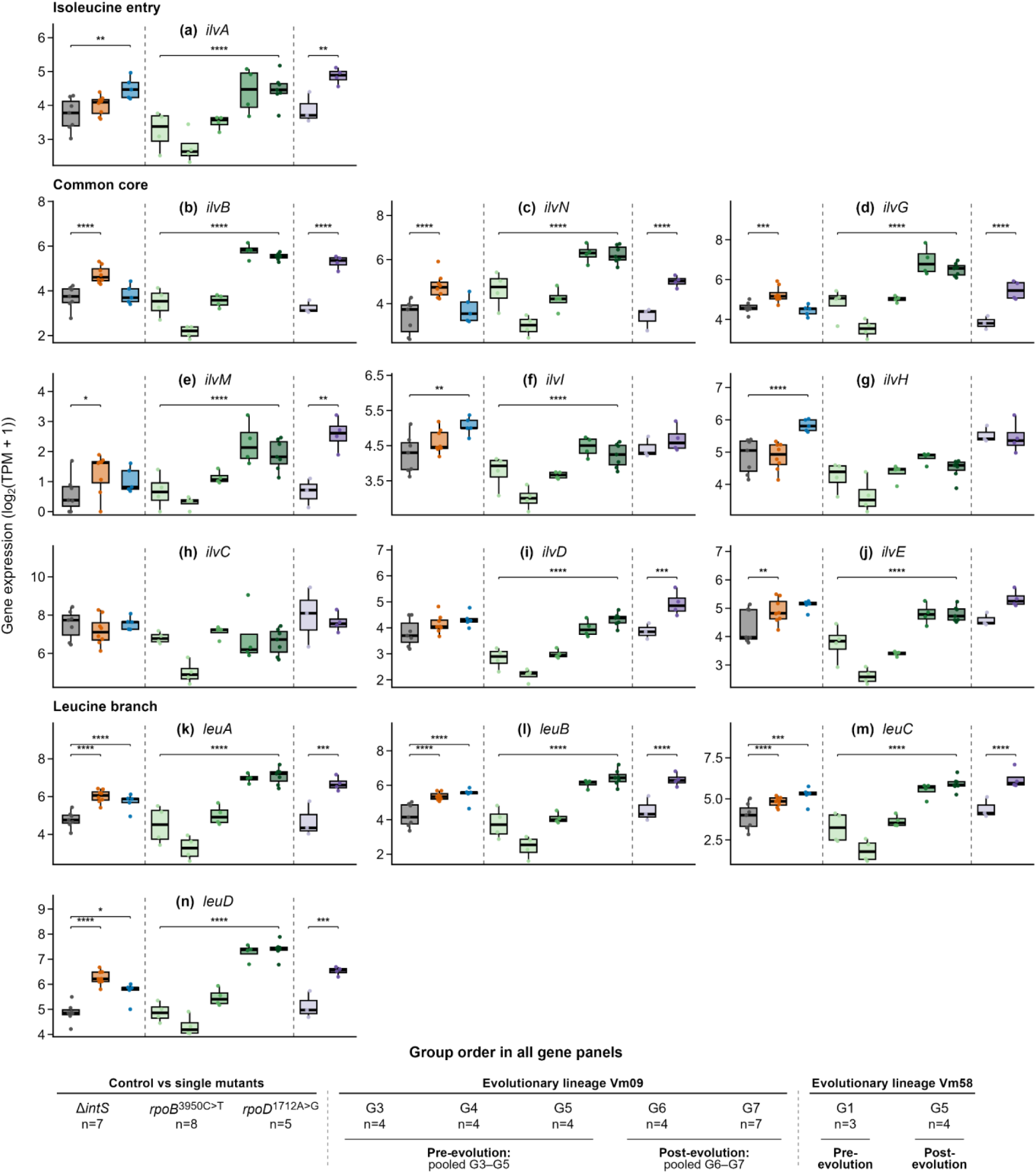
Expression of branched-chain amino acid biosynthesis genes across evolved lineages and reconstructed mutants. (**a–n**) Expression of *ilvA*, *ilvB*, *ilvN*, *ilvG*, *ilvM*, *ilvL*, *ilvH*, *ilvC*, *ilvD*, *ilvE*, *leuA*, *leuB*, *leuC* and *leuD*, respectively. In each panel, the left comparison is between Δ*intS* controls and reconstructed *rpoB*^3950C>T^ or *rpoD*^1712A>G^ mutants, the middle comparison is between Vm09 G3–G5 and G6–G7, and the right comparison is between Vm58 G1 and G5. Box plots show log_2_(TPM + 1); dots represent biological replicates, boxes show medians and IQRs, and whiskers extend to 1.5 × IQR. Statistical tests were performed on raw counts using DESeq2 Wald tests with Benjamini–Hochberg correction; log2 fold changes were shrunk using apeglm. Brackets denote prespecified comparisons with adjusted *P* < 0.05 and |shrunken log2 fold change| ≥ 0.5. \**P*_adj_ < 0.05, \*\**P*_adj_ < 0.01, \*\*\**P*_adj_ < 0.001 and \*\*\*\**P*_adj_ < 0.0001. Y-axis limits were scaled independently for each gene. Pathway assignments are shown in Fig. 5c.

**Fig. S10.**
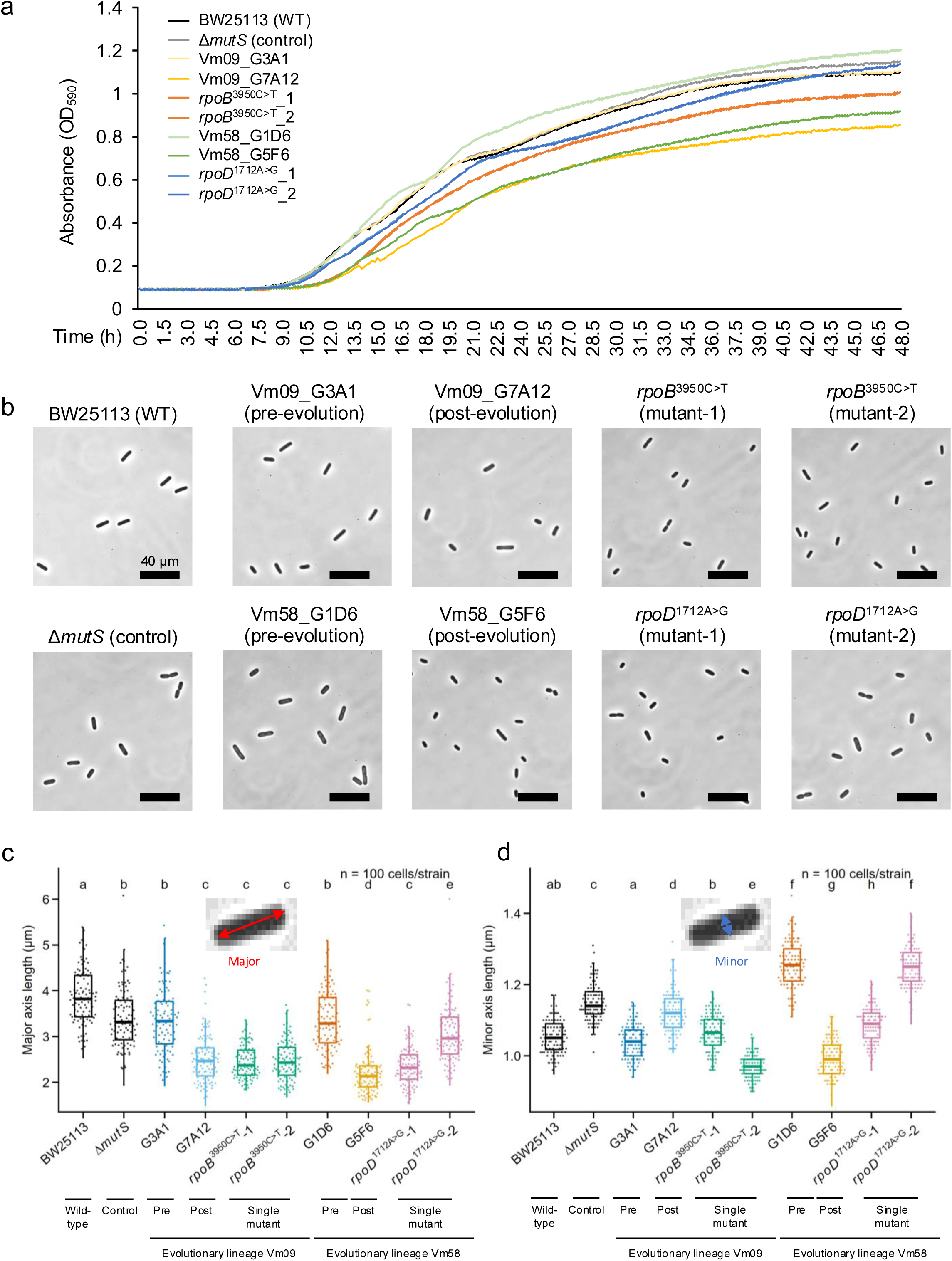
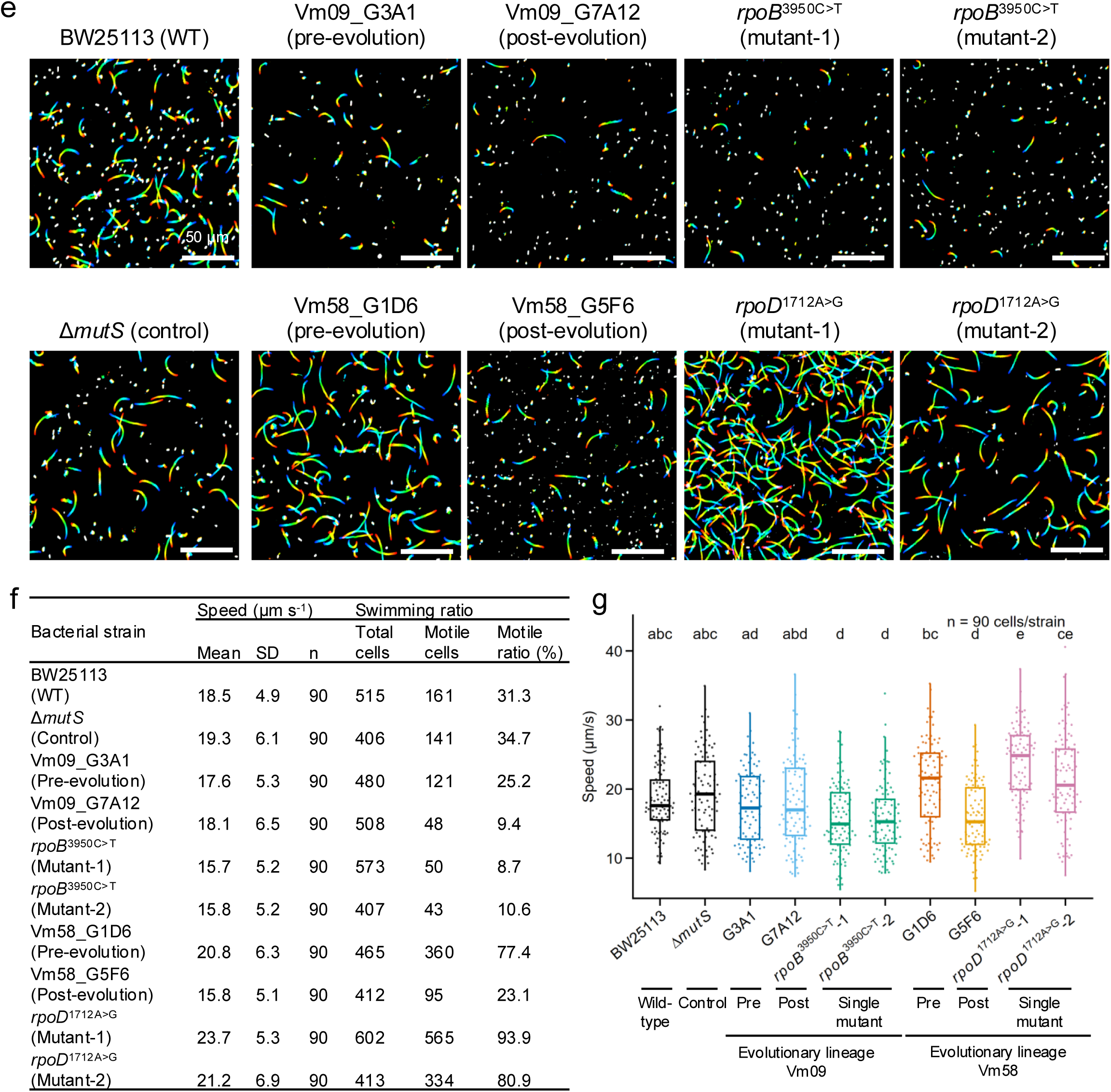
Growth, morphology and motility of evolved *E. coli* lineages, reconstructed *rpoB* and *rpoD* mutants and control strains. (**a**) Growth curves in liquid LB medium; lines show means of six replicate cultures. (**b**) Representative phase-contrast images. (**c,d**) Cell major-axis (**c**) and minor-axis (**d**) lengths. (**e**) One-second swimming trajectories visualized as rainbow plots. (**f,g**) Swimming speed and proportion of motile cells. In (**c,d,g**), different letters indicate significant differences (two-sided pairwise Wilcoxon rank-sum tests with Hommel correction; *P* < 0.05).

**Fig. S11.**
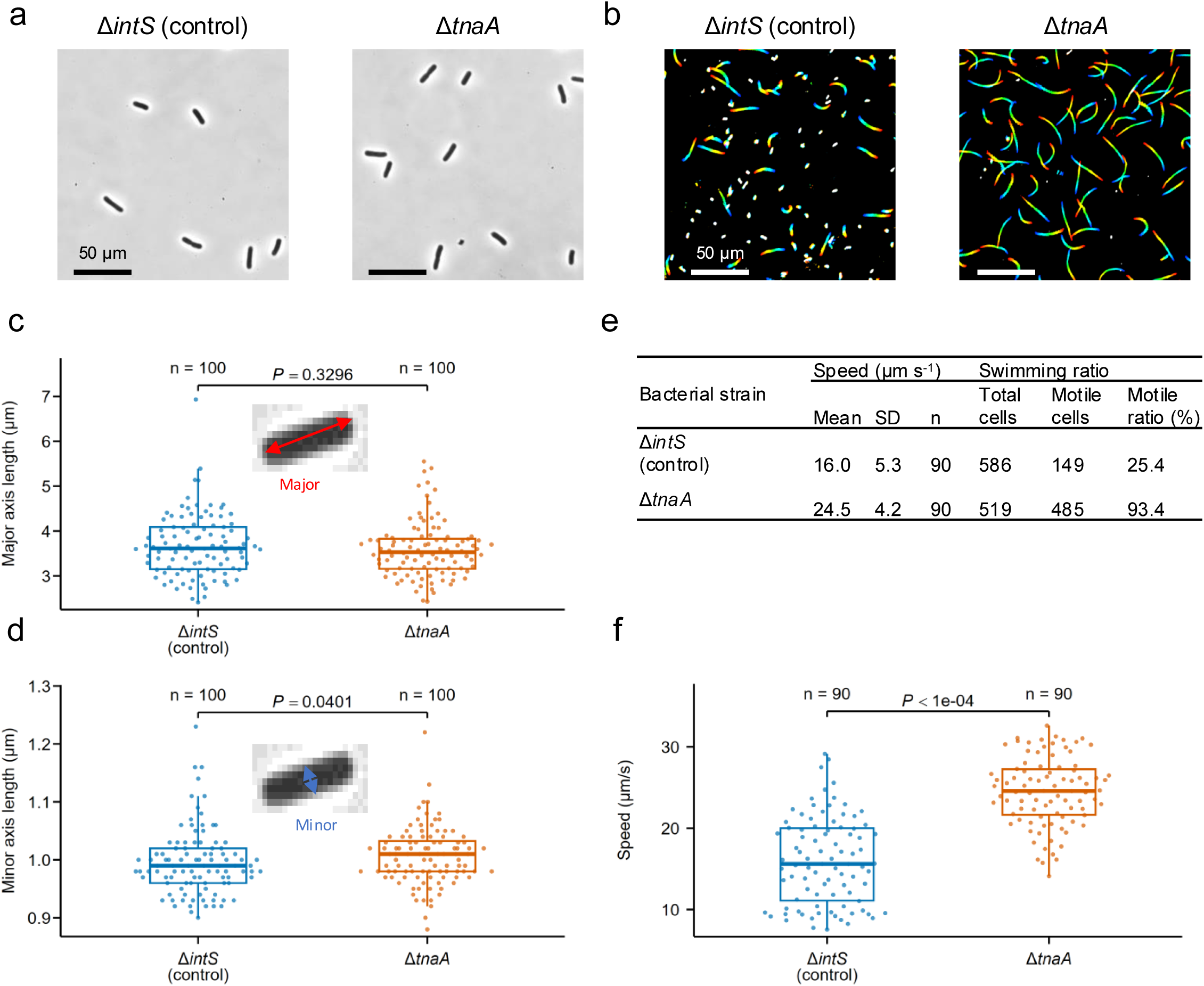
Morphology and motility of a mutualistic *E. coli* mutant Δ*tnaA* and a control strain Δ*intS*. (**a**) Representative phase-contrast images. (**b**) One-second swimming trajectories visualized as rainbow plots. (**c,d**) Cell major-axis and minor-axis lengths. (**e,f**) Swimming speed and proportion of motile cells. In (**c,d,f**), *P*-values of two-sided Wilcoxon rank–sum test are shown. In contrast to the disruptive mutants of the CCR global transcriptional regulator genes *cyaA* and *crp* showing reduced cell size and loss of flagellar motility^14^, the disruptive mutant of *tnaA* gene exhibited no reduced cell size and, unexpectedly, activated flagellar motility.

**Table S1.**
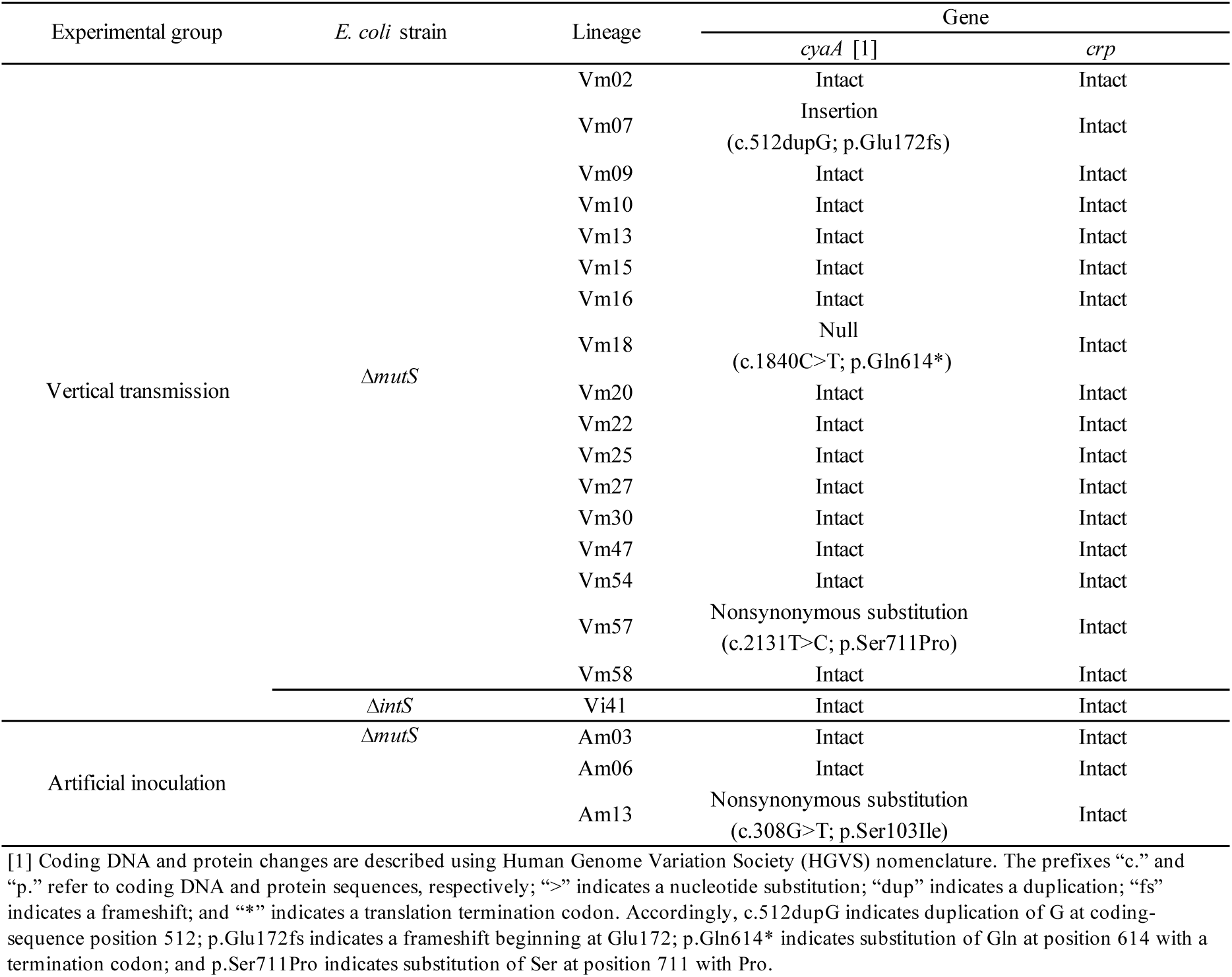
PCR amplification and Sanger sequencing of CCR component genes, *cyaA* and *crp*, in candidate evolutionary *E. coli* lineages.

**Table S2.**
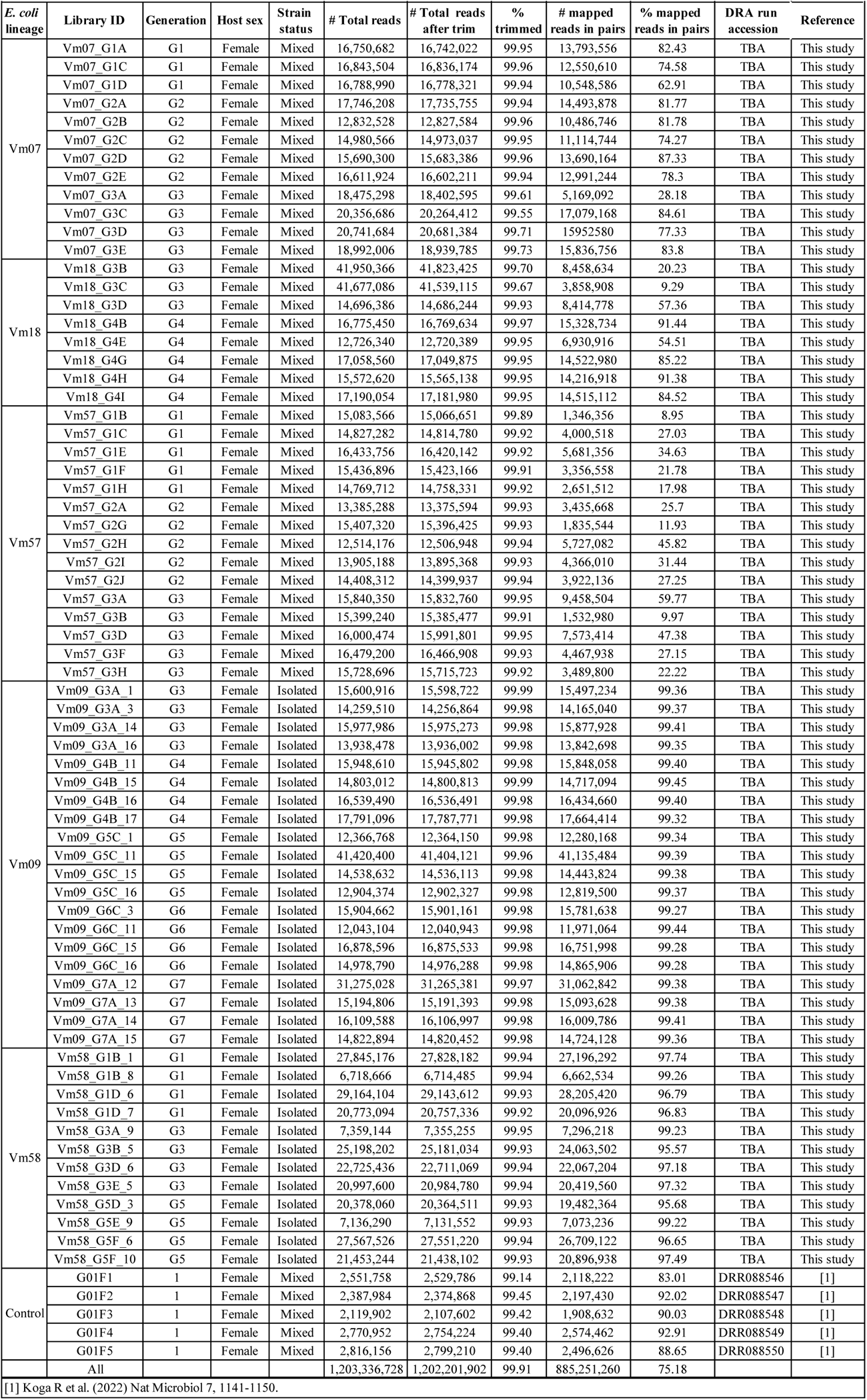
Basic statistics of DNA sequencing libraries.

**Table S3.**
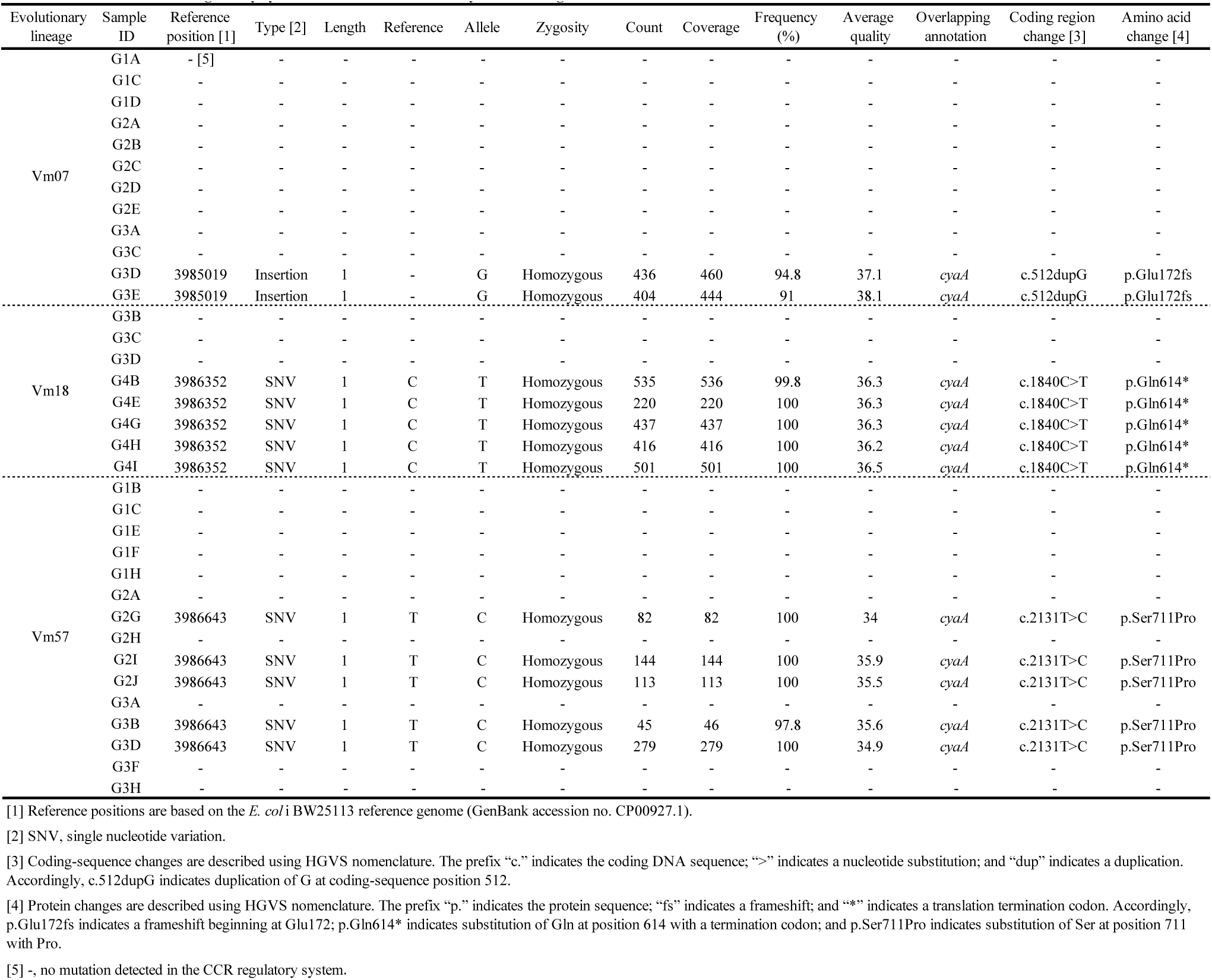
Genomics of CCR regulatory system mutations in evolutionary *E. coli* lineages.

**Table S4.** List of *E. coli* clones isolated from evolutionary *E. coli* lineage Vm09 for screening, infection experiments, genomics and transcriptomics.

| *E. coli* clones used for 1st screening |  |  |  |  |  | *E. coli* clones used for 2nd screening |  |  |  |  | DNA-seq | RNA-seq |  |
| --- | --- | --- | --- | --- | --- | --- | --- | --- | --- | --- | --- | --- | --- |
| No. | Control | Isolated evolved *E. coli* clone ID | Adult emergence rate (%) | Generation | Biological replicate | Wild type | Isolated evolved *E. coli* clone ID | Mean adult emergence rate (%) | Generation | Biological replicate | DNA-seq sample ID | RNA-seq sample ID | Biological replicate |
| 1 | *ΔintS* |  | 14.2 | G0 | 1 | *ΔintS* |  | 7.5 | G0 | 8 |  | *ΔintS* | 3 |
| 2 | *ΔintS* |  | 7.2 | G0 | 1 |  |  |  |  |  |  |  |  |
| 3 | *ΔintS* |  | 14.3 | G0 | 1 |  |  |  |  |  |  |  |  |
| 4 |  | Vm09_G1A_1 | 31.7 | G1 | 1 |  |  |  |  |  |  |  |  |
| 5 |  | Vm09_G1A_2 | 12.5 | G1 | 1 |  |  |  |  |  |  |  |  |
| 6 |  | Vm09_G1A_3 | 28.6 | G1 | 1 |  |  |  |  |  |  |  |  |
| 7 |  | Vm09_G1A_4 | 41.5 | G1 | 1 |  |  |  |  |  |  |  |  |
| 8 |  | Vm09_G1A_11 | 16.7 | G1 | 1 |  |  |  |  |  |  |  |  |
| 9 |  | Vm09_G1A_12 | 23.8 | G1 | 1 |  |  |  |  |  |  |  |  |
| 10 |  | Vm09_G1A_13 | 35.7 | G1 | 1 |  |  |  |  |  |  |  |  |
| 11 |  | Vm09_G1A_14 | 33.3 | G1 | 1 |  |  |  |  |  |  |  |  |
| 12 |  | Vm09_G1A_15 | 2.6 | G1 | 1 |  |  |  |  |  |  |  |  |
| 13 |  | Vm09_G1A_16 | 4.9 | G1 | 1 |  |  |  |  |  |  |  |  |
| 14 |  | Vm09_G1A_17 | 14.3 | G1 | 1 |  |  |  |  |  |  |  |  |
| 15 |  | Vm09_G2A_1 | 31.0 | G2 | 1 |  |  |  |  |  |  |  |  |
| 16 |  | Vm09_G2A_2 | 16.7 | G2 | 1 |  |  |  |  |  |  |  |  |
| 17 |  | Vm09_G2A_3 | 23.8 | G2 | 1 |  |  |  |  |  |  |  |  |
| 18 |  | Vm09_G2A_4 | 14.3 | G2 | 1 |  |  |  |  |  |  |  |  |
| 19 |  | Vm09_G2A_11 | 2.4 | G2 | 1 |  |  |  |  |  |  |  |  |
| 20 |  | Vm09_G2A_12 | 66.7 | G2 | 1 |  |  |  |  |  |  |  |  |
| 21 |  | Vm09_G2A_13 | 14.3 | G2 | 1 |  |  |  |  |  |  |  |  |
| 22 |  | Vm09_G2A_14 | 16.7 | G2 | 1 |  |  |  |  |  |  |  |  |
| 23 |  | Vm09_G2A_15 | 14.3 | G2 | 1 |  |  |  |  |  |  |  |  |
| 24 |  | Vm09_G2A_16 | 0.0 | G2 | 1 |  |  |  |  |  |  |  |  |
| 25 |  | Vm09_G2A_17 | 31.0 | G2 | 1 |  |  |  |  |  |  |  |  |
| 26 |  | Vm09_G3A_1 | 20.0 | G3 | 1 |  | Vm09_G3A_1 | 24.3 | G3 | 6 | Vm09_G3A_1 |  |  |
| 27 |  | Vm09_G3A_2 | 28.6 | G3 | 1 |  |  |  |  |  |  |  |  |
| 28 |  | Vm09_G3A_3 | 17.1 | G3 | 1 |  | Vm09_G3A_3 | 21.5 | G3 | 6 | Vm09_G3A_3 |  |  |
| 29 |  | Vm09_G3A_11 | 61.9 | G3 | 1 |  |  |  |  |  |  |  |  |
| 30 |  | Vm09_G3A_12 | 33.3 | G3 | 1 |  |  |  |  |  |  |  |  |
| 31 |  | Vm09_G3A_13 | 59.5 | G3 | 1 |  |  |  |  |  |  |  |  |
| 32 |  | Vm09_G3A_14 | 4.8 | G3 | 1 |  | Vm09_G3A_14 | 28.4 | G3 | 6 | Vm09_G3A_14 |  |  |
| 33 |  | Vm09_G3A_15 | 36.6 | G3 | 1 |  |  |  |  |  |  |  |  |
| 34 |  | Vm09_G3A_16 | 9.8 | G3 | 1 |  | Vm09_G3A_16 | 26.3 | G3 | 6 | Vm09_G3A_16 | Vm09_G3A_16 | 4 |
| 35 |  | Vm09_G3A_17 | 25.0 | G3 | 1 |  |  |  |  |  |  |  |  |
| 36 |  | Vm09_G4B_1 | 36.6 | G4 | 1 |  |  |  |  |  |  |  |  |
| 37 |  | Vm09_G4B_2 | 34.2 | G4 | 1 |  |  |  |  |  |  |  |  |
| 38 |  | Vm09_G4B_3 | 22.0 | G4 | 1 |  |  |  |  |  |  |  |  |
| 39 |  | Vm09_G4B_11 | 4.8 | G4 | 1 |  | Vm09_G4B_11 | 17.7 | G4 | 7 | Vm09_G4B_11 |  |  |
| 40 |  | Vm09_G4B_12 | 38.1 | G4 | 1 |  |  |  |  |  |  |  |  |
| 41 |  | Vm09_G4B_13 | 28.6 | G4 | 1 |  |  |  |  |  |  |  |  |
| 42 |  | Vm09_G4B_14 | 11.9 | G4 | 1 |  |  |  |  |  |  |  |  |
| 43 |  | Vm09_G4B_15 | 7.7 | G4 | 1 |  | Vm09_G4B_15 | 25.2 | G4 | 7 | Vm09_G4B_15 | Vm09_G4B_15 | 4 |
| 44 |  | Vm09_G4B_16 | 2.5 | G4 | 1 |  | Vm09_G4B_16 | 13.9 | G4 | 7 | Vm09_G4B_16 |  |  |
| 45 |  | Vm09_G4B_17 | 7.3 | G4 | 1 |  | Vm09_G4B_17 | 24.1 | G4 | 7 | Vm09_G4B_17 |  |  |
| 46 |  | Vm09_G5C_1 | 63.4 | G5 | 1 |  | Vm09_G5C_1 | 36.6 | G5 | 8 | Vm09_G5C_1 |  |  |
| 47 |  | Vm09_G5C_2 | 31.0 | G5 | 1 |  |  |  |  |  |  |  |  |
| 48 |  | Vm09_G5C_3 | 34.1 | G5 | 1 |  |  |  |  |  |  |  |  |
| 49 |  | Vm09_G5C_11 | 61.9 | G5 | 1 |  | Vm09_G5C_11 | 26.9 | G5 | 8 | Vm09_G5C_11 |  |  |
| 50 |  | Vm09_G5C_12 | 57.1 | G5 | 1 |  |  |  |  |  |  |  |  |
| 51 |  | Vm09_G5C_13 | 50.0 | G5 | 1 |  |  |  |  |  |  |  |  |
| 52 |  | Vm09_G5C_14 | 50.0 | G5 | 1 |  |  |  |  |  |  |  |  |
| 53 |  | Vm09_G5C_15 | 61.0 | G5 | 1 |  | Vm09_G5C_15 | 34.9 | G5 | 8 | Vm09_G5C_15 |  |  |
| 54 |  | Vm09_G5C_16 | 71.4 | G5 | 1 |  | Vm09_G5C_16 | 43.1 | G5 | 8 | Vm09_G5C_16 | Vm09_G5C_16 | 4 |
| 55 |  | Vm09_G6C_1 | 69.0 | G6 | 1 |  |  |  |  |  |  |  |  |
| 56 |  | Vm09_G6C_2 | 0.0 | G6 | 1 |  |  |  |  |  |  |  |  |
| 57 |  | Vm09_G6C_3 | 80.5 | G6 | 1 |  | Vm09_G6C_3 | 56.2 | G6 | 8 | Vm09_G6C_3 |  |  |
| 58 |  | Vm09_G6C_11 | 81.0 | G6 | 1 |  | Vm09_G6C_11 | 47.3 | G6 | 8 | Vm09_G6C_11 | Vm09_G6C_11 | 4 |
| 59 |  | Vm09_G6C_12 | 71.4 | G6 | 1 |  |  |  |  |  |  |  |  |
| 60 |  | Vm09_G6C_13 | 54.8 | G6 | 1 |  |  |  |  |  |  |  |  |
| 61 |  | Vm09_G6C_14 | 36.6 | G6 | 1 |  |  |  |  |  |  |  |  |
| 62 |  | Vm09_G6C_15 | 75.0 | G6 | 1 |  | Vm09_G6C_15 | 43.9 | G6 | 8 | Vm09_G6C_15 |  |  |
| 63 |  | Vm09_G6C_16 | 75.0 | G6 | 1 |  | Vm09_G6C_16 | 46.4 | G6 | 8 | Vm09_G6C_16 |  |  |
| 64 |  | Vm09_G7A_1 | 2.6 | G7 | 1 |  |  |  |  |  |  |  |  |
| 65 |  | Vm09_G7A_2 | 42.1 | G7 | 1 |  |  |  |  |  |  |  |  |
| 66 |  | Vm09_G7A_3 | 42.9 | G7 | 1 |  |  |  |  |  |  |  |  |
| 67 |  | Vm09_G7A_11 | 64.3 | G7 | 1 |  |  |  |  |  |  |  |  |
| 68 |  | Vm09_G7A_12 | 76.2 | G7 | 1 |  | Vm09_G7A_12 | 56.4 | G7 | 8 | Vm09_G7A_12 | Vm09_G7A_12 | 3 |
| 69 |  | Vm09_G7A_13 | 76.2 | G7 | 1 |  | Vm09_G7A_13 | 47.9 | G7 | 8 | Vm09_G7A_13 | Vm09_G7A_13 | 4 |
| 70 |  | Vm09_G7A_14 | 66.7 | G7 | 1 |  | Vm09_G7A_14 | 49.8 | G7 | 8 | Vm09_G7A_14 |  |  |
| 71 |  | Vm09_G7A_15 | 75.6 | G7 | 1 |  | Vm09_G7A_15 | 59.5 | G7 | 8 | Vm09_G7A_15 |  |  |
| 72 |  | Vm09_G7A_16 | 61.9 | G7 | 1 |  |  |  |  |  |  |  |  |

**Table S5.**
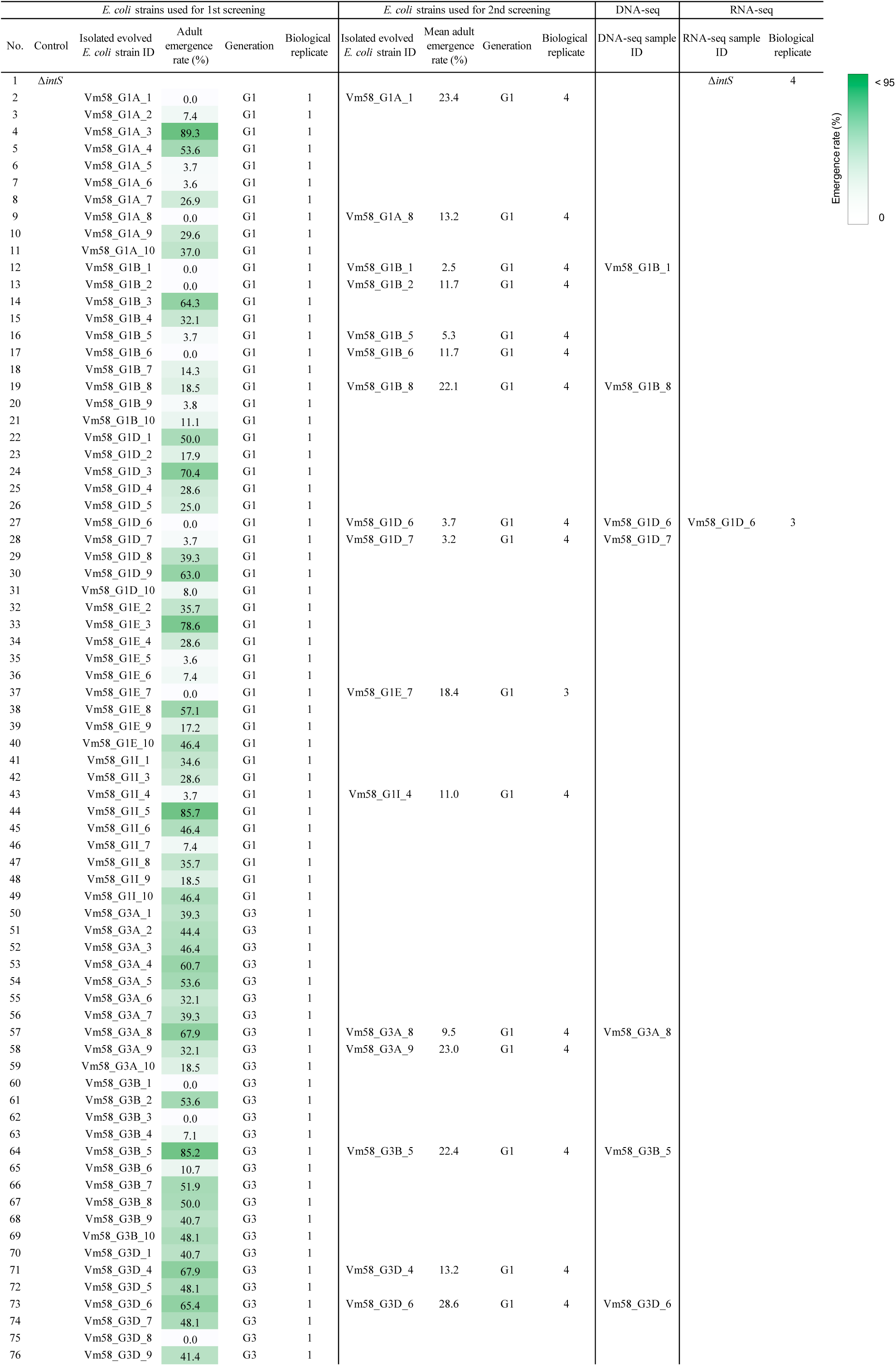
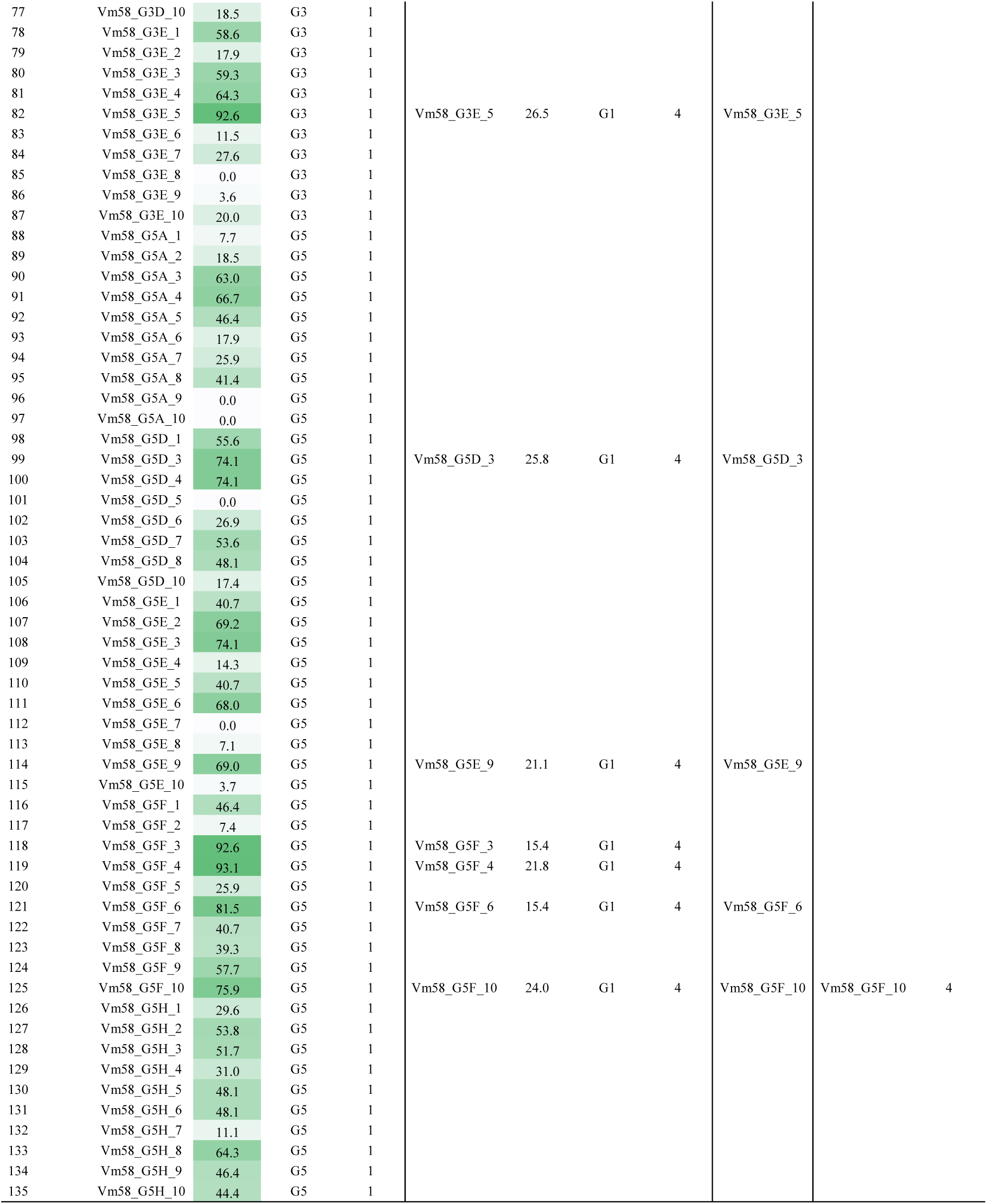
List of *E. coli* clones isolated from evolutionary *E. coli* lineage Vm58 for screening, infection experiments, genomics and transcriptomics.

**Table S6.** Basic statistics of RNA sequencing libraries.

| *E. coli*<br>lineage | Status | Library ID | Generation | Sex | Chemistry | # Total reads | # Total reads<br>after trim | % QC<br>passed reads | # mapped<br>reads in pairs | % mapped<br>in pairs | DRA run<br>accession |
| --- | --- | --- | --- | --- | --- | --- | --- | --- | --- | --- | --- |
| Vm09 | Isolated | Vm09_G3A_16F1 | 3 | Female | TruSeq RNA | 81,056,888 | 81,021,696 | 99.96 | 13,785,756 | 17.02 | TBA |
|  | Isolated | Vm09_G3A_16F2 | 3 | Female | TruSeq RNA | 66,227,210 | 66,193,287 | 99.95 | 10,422,278 | 15.75 | TBA |
|  | Isolated | Vm09_G3A_16F3 | 3 | Female | TruSeq RNA | 53,384,312 | 53,361,376 | 99.96 | 7,040,042 | 13.20 | TBA |
|  | Isolated | Vm09_G3A_16F4 | 3 | Female | TruSeq RNA | 40,974,638 | 40,947,582 | 99.93 | 5,762,746 | 14.08 | TBA |
|  | Isolated | Vm09_G4B_15F1 | 4 | Female | TruSeq RNA | 50,462,696 | 50,439,832 | 99.95 | 7,068,372 | 14.02 | TBA |
|  | Isolated | Vm09_G4B_15F2 | 4 | Female | TruSeq RNA | 539,737,370 | 539,433,638 | 99.94 | 121,495,080 | 22.53 | TBA |
|  | Isolated | Vm09_G4B_15F3 | 4 | Female | TruSeq RNA | 463,558,374 | 463,347,195 | 99.95 | 127,526,452 | 27.53 | TBA |
|  | Isolated | Vm09_G4B_15F4 | 4 | Female | TruSeq RNA | 260,211,052 | 260,123,755 | 99.97 | 39,031,054 | 15.01 | TBA |
|  | Isolated | Vm09_G5C_16F1 | 5 | Female | TruSeq RNA | 122,390,918 | 122,331,942 | 99.95 | 30,152,338 | 24.66 | TBA |
|  | Isolated | Vm09_G5C_16F2 | 5 | Female | TruSeq RNA | 62,754,156 | 62,722,230 | 99.95 | 15,089,032 | 30.60 | TBA |
|  | Isolated | Vm09_G5C_16F3 | 5 | Female | TruSeq RNA | 53,969,894 | 53,935,115 | 99.94 | 15,275,640 | 28.33 | TBA |
|  | Isolated | Vm09_G5C_16F4 | 5 | Female | TruSeq RNA | 62,754,156 | 62,722,230 | 99.95 | 14,072,288 | 22.44 | TBA |
|  | Isolated | Vm09_G6C_11F1 | 6 | Female | TruSeq RNA | 73,629,384 | 73,601,004 | 99.96 | 19,286,600 | 26.21 | TBA |
|  | Isolated | Vm09_G6C_11F2 | 6 | Female | TruSeq RNA | 47,927,290 | 47,905,862 | 99.96 | 9,052,632 | 18.90 | TBA |
|  | Isolated | Vm09_G6C_11F3 | 6 | Female | TruSeq RNA | 71,112,142 | 71,067,858 | 99.94 | 20,231,470 | 28.47 | TBA |
|  | Isolated | Vm09_G6C_11F4 | 6 | Female | TruSeq RNA | 50,410,988 | 50,385,293 | 99.95 | 12,486,538 | 24.79 | TBA |
|  | Isolated | Vm09_G7A_12F1 | 7 | Female | TruSeq RNA | 83,243,506 | 83,200,179 | 99.95 | 13,974,428 | 16.80 | TBA |
|  | Isolated | Vm09_G7A_12F2 | 7 | Female | TruSeq RNA | 78,246,038 | 78,186,135 | 99.92 | 18,731,322 | 23.96 | TBA |
|  | Isolated | Vm09_G7A_12F3 | 7 | Female | TruSeq RNA | 69,266,762 | 69,179,846 | 99.87 | 11,554,446 | 16.71 | TBA |
|  | Isolated | Vm09_G7A_13F1 | 7 | Female | TruSeq RNA | 174,044,684 | 173,997,617 | 99.97 | 34,116,678 | 19.61 | TBA |
|  | Isolated | Vm09_G7A_13F2 | 7 | Female | TruSeq RNA | 187,611,926 | 187,544,454 | 99.96 | 38,553,074 | 20.56 | TBA |
|  | Isolated | Vm09_G7A_13F3 | 7 | Female | TruSeq RNA | 194,929,440 | 194,857,079 | 99.96 | 42,158,092 | 21.64 | TBA |
|  | Isolated | Vm09_G7A_13F4 | 7 | Female | TruSeq RNA | 162,848,332 | 162,768,655 | 99.95 | 25,531,390 | 15.69 | TBA |
|  | Single mutant | *rpoB*(2)<sup>3950C>T</sup>F1 | - | Female | TruSeq RNA | 191,174,798 | 191,083,769 | 99.95 | 58,254,964 | 30.49 | TBA |
|  | Single mutant | *rpoB*(2)<sup>3950C>T</sup>F2 | - | Female | TruSeq RNA | 190,388,774 | 190,341,971 | 99.98 | 41,042,852 | 21.56 | TBA |
|  | Single mutant | *rpoB*(2)<sup>3950C>T</sup>F3 | - | Female | TruSeq RNA | 138,894,542 | 138,864,754 | 99.98 | 9,852,090 | 7.10 | TBA |
|  | Single mutant | *rpoB*(2)<sup>3950C>T</sup>F4 | - | Female | TruSeq RNA | 183,988,654 | 183,914,118 | 99.96 | 42,191,046 | 22.94 | TBA |
|  | Single mutant | *rpoB*(3)<sup>3950C>T</sup>F1 | - | Female | TruSeq RNA | 200,852,548 | 200,786,669 | 99.97 | 54,247,826 | 27.02 | TBA |
|  | Single mutant | *rpoB*(3)<sup>3950C>T</sup>F2 | - | Female | TruSeq RNA | 157,244,568 | 157,166,198 | 99.95 | 19,470,982 | 12.39 | TBA |
|  | Single mutant | *rpoB*(3)<sup>3950C>T</sup>F3 | - | Female | TruSeq RNA | 146,398,494 | 131,918,740 | 90.11 | 14,110,042 | 9.64 | TBA |
|  | Single mutant | *rpoB*(3)<sup>3950C>T</sup>F4 | - | Female | TruSeq RNA | 170,467,196 | 170,406,808 | 99.96 | 46,688,722 | 27.40 | TBA |
| | Control | BW25113 $\Delta intS$_F1 | - | Female | TruSeq RNA | 209,844,068 | 209,784,839 | 99.97 | 56,086,822 | 26.74 | TBA |
| | Control | BW25113 $\Delta intS$_F2 | - | Female | TruSeq RNA | 215,682,070 | 215,602,880 | 99.96 | 52,930,222 | 24.55 | TBA |
| | Control | BW25113 $\Delta intS$_F3 | - | Female | TruSeq RNA | 156,878,782 | 156,840,808 | 99.98 | 26,058,738 | 16.62 | TBA |
| Vm58 | Isolated | Vm58_G1D_6F1 | 1 | Female | TruSeq RNA | 154,763,486 | 153,380,797 | 99.11 | 30,003,330 | 19.63 | TBA |
|  | Isolated | Vm58_G1D_6F2 | 1 | Female | TruSeq RNA | 235,941,154 | 233,771,246 | 99.08 | 53,909,842 | 23.14 | TBA |
|  | Isolated | Vm58_G1D_6F3 | 1 | Female | TruSeq RNA | 272,219,596 | 268,920,360 | 98.79 | 71,852,362 | 26.81 | TBA |
|  | Isolated | Vm58_G5F_10F1 | 5 | Female | TruSeq RNA | 302,188,626 | 299,241,615 | 99.02 | 39,702,274 | 13.31 | TBA |
|  | Isolated | Vm58_G5F_10F2 | 5 | Female | TruSeq RNA | 277,701,186 | 275,361,517 | 99.16 | 27,866,628 | 10.15 | TBA |
|  | Isolated | Vm58_G5F_10F3 | 5 | Female | TruSeq RNA | 170,703,084 | 167,373,782 | 98.05 | 18,585,816 | 11.15 | TBA |
|  | Isolated | Vm58_G5F_10F4 | 5 | Female | TruSeq RNA | 54,676,440 | 54,191,646 | 99.11 | 4,217,008 | 7.80 | TBA |
|  | Single mutant | *rpoD*(16)<sup>1712A>G</sup>F1 | - | Female | TruSeq RNA | 237,727,182 | 235,710,334 | 99.15 | 25,146,458 | 10.70 | TBA |
|  | Single mutant | *rpoD*(16)<sup>1712A>G</sup>F2 | - | Female | TruSeq RNA | 355,435,564 | 352,969,091 | 99.31 | 45,864,372 | 13.03 | TBA |
|  | Single mutant | *rpoD*(16)<sup>1712A>G</sup>F3 | - | Female | TruSeq RNA | 292,071,334 | 289,912,678 | 99.26 | 30,591,028 | 10.58 | TBA |
|  | Single mutant | *rpoD*(16)<sup>1712A>G</sup>F4 | - | Female | TruSeq RNA | 273,097,022 | 271,190,820 | 99.30 | 23,680,572 | 8.76 | TBA |
|  | Single mutant | *rpoD*(16)<sup>1712A>G</sup>F5 | - | Female | TruSeq RNA | 194,688,800 | 192,832,740 | 99.05 | 27,720,830 | 14.42 | TBA |
| | Control | BW25113 $\Delta intS$_F5 | - | Female | TruSeq RNA | 67,516,012 | 67,145,336 | 99.45 | 17,317,250 | 25.86 | TBA |
| | Control | BW25113 $\Delta intS$_F6 | - | Female | TruSeq RNA | 118,911,342 | 118,162,229 | 99.37 | 33,880,876 | 28.75 | TBA |
| | Control | BW25113 $\Delta intS$_F7 | - | Female | TruSeq RNA | 142,901,676 | 142,026,746 | 99.39 | 33,883,570 | 23.93 | TBA |
| | Control | BW25113 $\Delta intS$_F8 | - | Female | TruSeq RNA | 231,563,592 | 229,549,693 | 99.13 | 50,840,174 | 22.22 | TBA |
| All |  |  |  |  |  | 8,394,672,746 | 8,347,726,044 | 99.44 | 1,608,394,444 | 18.75 |  |

**Table S7.**
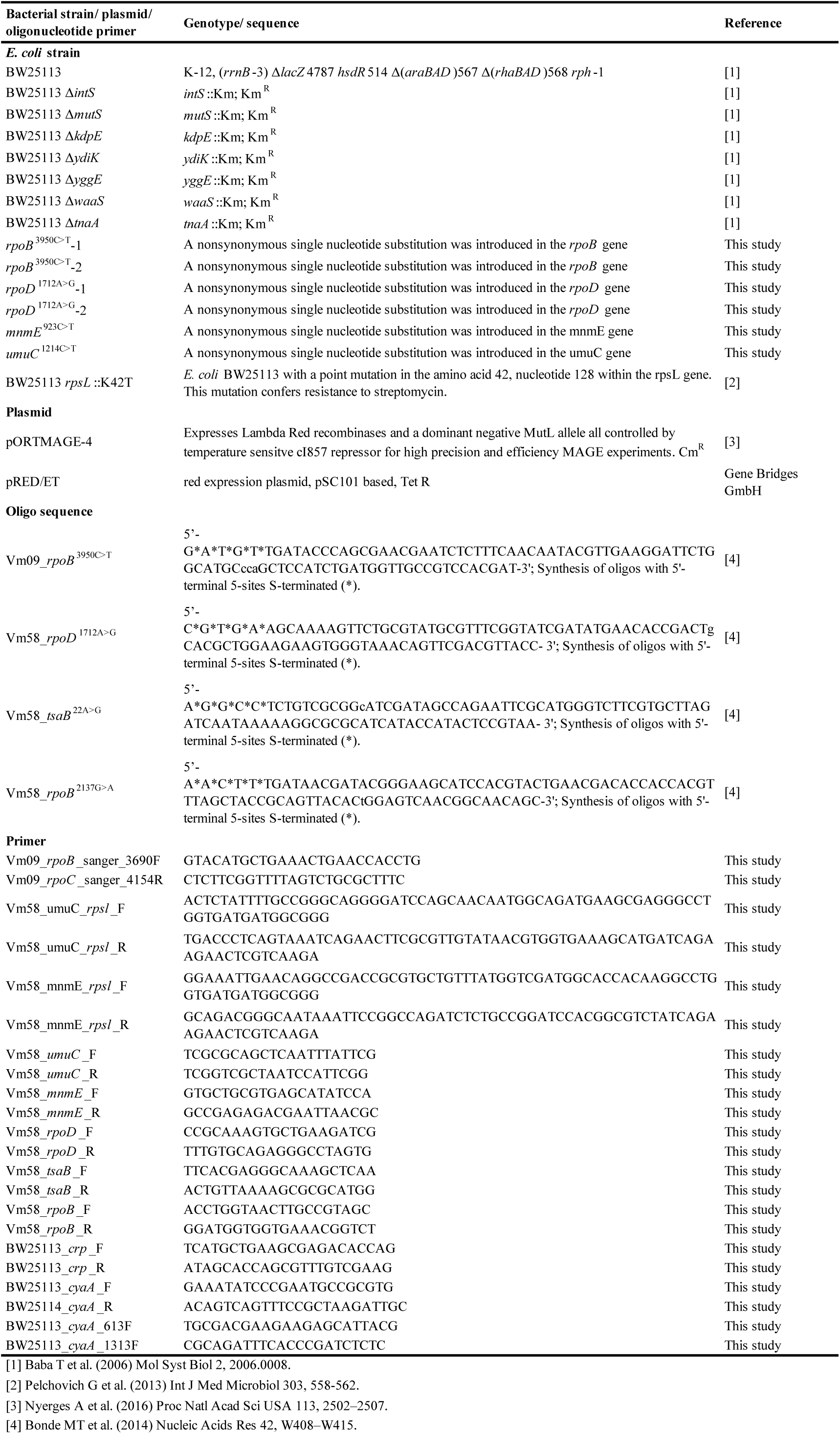
Bacterial strains, plasmids and oligonucleotide sequences used in this study.

